# The tRNA dihydrouridine synthase DusA has a distinct mechanism in optimizing tRNAs for translation

**DOI:** 10.1101/2025.08.28.672980

**Authors:** Sarah K. Schultz, Nadia Hossain, Lauren Barnes, Kristin S. Koutmou, Ute Kothe

## Abstract

Dihydrouridine (D) is one of the most highly conserved RNA modifications across all domains of life. D20 within the tRNA D loop is particularly conserved and is formed by DusA in *Escherichia coli.* However, the mechanisms and cellular functions of DusA and D20 remain poorly understood. Here, we characterize DusA’s role in tRNA binding, cofactor oxidation, and modification activity, along with its impact on tRNA maturation and translation. We find that DusA binds tRNA via a two-step mechanism involving a local structural rearrangement and exhibits a higher affinity for previously modified tRNA compared to unmodified tRNA. Unlike the T arm modifying enzymes TrmA and TruB, DusA does not broadly increase cellular aminoacylation for all tRNAs but enhances the charging of specific tRNA species. Despite limited alterations in overall tRNA charging and abundance in cells lacking DusA, DusA selectively improves translation at several specific codons, suggesting a direct contribution for dihydrouridine to the function of certain tRNAs on the ribosome. In conclusion, our findings suggest DusA acts non-redundantly with and complementary to TrmA and TruB in fine-tuning protein synthesis.

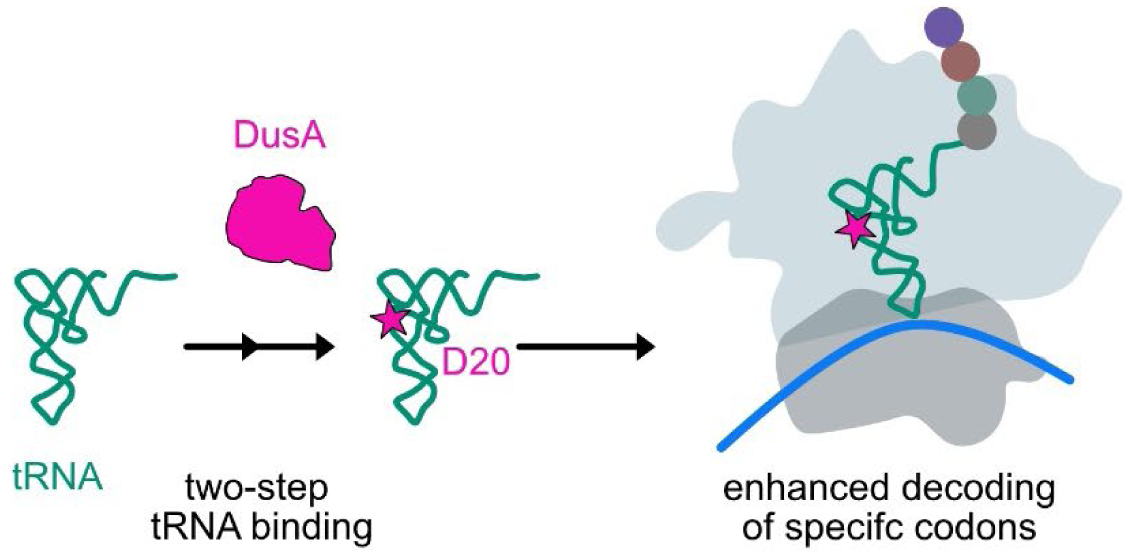

## Introduction

All transfer RNAs (tRNAs) are extensively and diversely modified (1). The RNA modifying enzymes that introduce tRNA modifications play a variety of roles in tRNA maturation and function (2). Across all domains of life, almost every tRNA features at least one dihydrouridine (D) modification, found most often within the eponymous tRNA D loop (3–5). Whereas most RNA modifications stabilize the tRNA structure, the unique non-planar nucleobase of dihydrouridine cannot participate in stabilizing base stacking interactions and primarily adopts the flexible C2ʹ endo ribose conformation (Figure 1A) (4–6). As such, dihydrouridine provides local flexibility within the tRNA structure, which may facilitate tertiary base pairing in the tRNA elbow, thereby stabilizing the overall tRNA structure (7,8). Supporting a functional role for dihydrouridine in increasing tRNA flexibility, tRNAs from psychrophilic organisms tend to contain more dihydrouridine than tRNAs from mesophiles and thermophiles (9,10).

**Figure 1.**
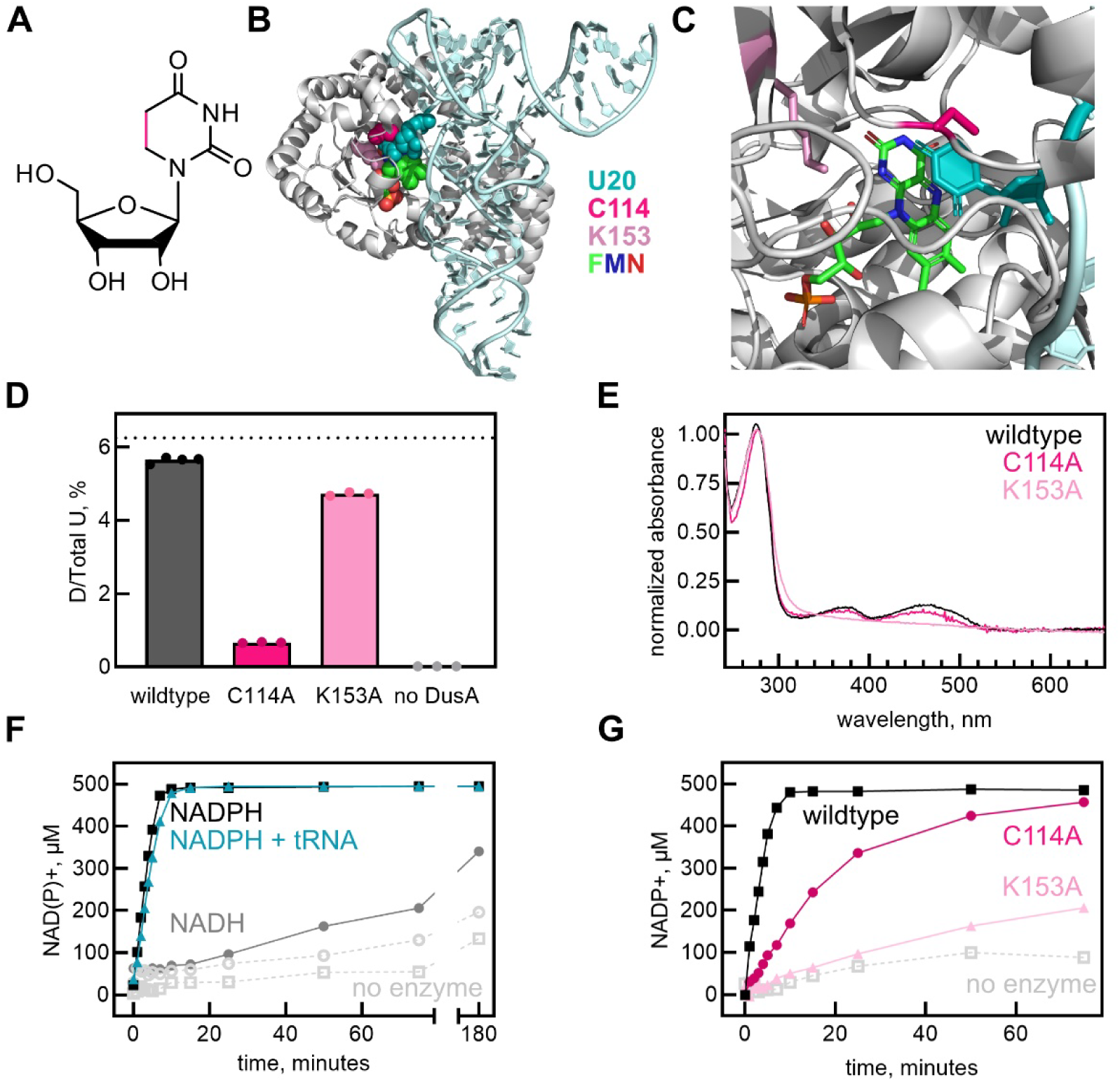
Characterization of *E. coli* DusA activity. **(A)** Structure of dihydrouridine, with the reduced C5-C6 bond as compared to uridine colored in pink. **(B)** Structure of *T. thermophilus* Dus (*Tth*Dus) in complex with tRNA^Phe^ and FMN (PDB 3B0V) (32). *Tth*Dus is grey, tRNA^Phe^ is light teal, and FMN is shown as spheres and coloured by atom (green: carbon, blue: nitrogen, red: oxygen). Residues C93 (equivalent to *E. coli* DusA C114) and K132 (equivalent to *E. coli* DusA K153) are shown as spheres, coloured in dark pink and light pink, respectively (residue numbering in figure is according to *E. coli* numbering). The target base, U20, is shown as spheres and coloured teal. **(C)** Zoomed in view of the *Tth*Dus active site, with residues coloured as in panel B, but shown as sticks. **(D)** Dihydrouridine formation following one hour incubation of equimolar concentrations of *in vitro* transcribed tRNA^Phe^ and purified DusA enzyme or no enzyme in the presence of FMN and NADPH (n=3 independent modification reactions) determined by mass spectrometry. *E. coli* tRNA^Phe^ contains 16 U nucleotides; therefore, 100% modification at one site (U20) corresponds to 6.25% D/total U (dashed line). **(E)** Absorbance spectra for purified *E. coli* DusA wildtype (black), and C114A (dark pink) and K153A (light pink) variants. **(F)** Oxidation of 500 µM NADPH (black squares) or NADH (grey circles) by 2 µM DusA. No enzyme controls for NADPH and NADH are shown as light grey squares and circles with dashed lines, respectively. **(G)** Oxidation of 500 µM NADPH to form NADP+ by 2 µM DusA wildtype (black), C114A (dark pink), and K153A (light pink). Table 2 lists initial velocities for each reaction in panels F and G.

It is speculated that all dihydrouridine synthases (Dus) arose from duplications of a common ancestral Dus (11,12). Bacteria have three major families of tRNA dihydrouridine synthases, DusA, DusB, and DusC (13), in addition to a recently discovered rRNA dihydrouridine synthase, RdsA (14). In *Escherichia coli*, DusA, B, and C have non-overlapping specificities, with DusA modifying U20 and U20a, DusB modifying U17, and DusC modifying D16 (13). DusB is regarded to be the ancestral protein and DusA and DusC are thought to have arisen from gene duplication (11,12). However, dihydrouridine synthase evolution in prokaryotes is complex; for example, Firmicutes harbour up to three DusB subgroups (DusB1-3), wherein DusB1 homologues can have multisite specificity that varies between species (15,16). Eukaryotes have four families of dihydrouridine synthases with non-overlapping specificities, Dus1-4, wherein Dus1 modifies both U16 and U17, Dus2 modifies U20, Dus3 modifies U47 within the tRNA variable loop, and Dus4 modifies U20a and U20b (17). Finally, archaeal Dus constitutes a separate family of dihydrouridine synthases that are relatively diverged from each other and not well-characterized to date (11).

DusA modifies position 20 and/or 20a in over 35 of the 46 *E. coli* tRNAs (18), thereby contributing about half of the bulk cellular tRNA dihydrouridine content (13). Like other tRNA modifying enzymes, the *dusA* gene can be deleted from *E. coli* and, even in combination with additional deletion of *dusB* and *dusC* genes, no growth defects are present for the triple knockout strain grown in ideal conditions (13) or for deletion of the *dusA* gene from *Acinetobacter baumannii* in several stress conditions (19). Similarly, no significant growth phenotype was reported for yeast lacking *dus1, dus2, dus3,* or *dus4* in solid and liquid media at various temperatures (17). Since the *dusA* gene and D20/D20a modifications are highly conserved across Proteobacteria (11,12) and DusB orthologs have been identified to instead introduce D20/D20a modifications in several Gram-positive species that lack DusA (15,16), tRNA U20/U20a dihydrouridylation is likely to contribute to cellular fitness, as has been shown previously for other tRNA modifying enzymes (20). In a screen of several tRNA modifying enzymes, competition experiments and transposon insertion sequencing revealed that *dusA* inactivation is actually beneficial to *E. coli* and *Vibrio cholerae* fitness when grown in sub-minimal inhibitory concentrations of the aminoglycoside antibiotic tobramycin (21). Similar enhancements in cell growth under certain stress conditions have also been described for other tRNA modifying enzyme knockouts including *E. coli trmA* and its homolog yeast *trm2* (22), but this increase in sensitivity to stressors does not explain why these genes are so highly conserved. Interestingly, the *dusA* gene has been shown to serve as an integration site for prophages and genomic islands of diverse functions in over 200 sequenced Proteobacterial organisms (19,23). Although genetic element integration disrupts *dusA*, in investigated cases, a new promoter and restored reading frame is provided, allowing for continued expression of DusA protein (23,24).

In humans, dihydrouridine has been long known to be overabundant in tRNAs from tumor cells compared to healthy tissue (25), and the U20-modifying enzyme hDus2 has been shown to be overexpressed in clinical lung cancer samples and non-small cell lung cancer cell lines (26). Indeed, hDus2 is necessary for survival and growth of lung cancer cells, and hDus2 expression inversely correlates with patient survival time (26). The mechanism underlying the involvement of hDus2 and cancer progression remains unclear. Interestingly, hDus2 has been shown to interact with glutamyl-prolyl tRNA synthetase and protein kinase R, which may have biological significance in cancer cells (26,27). In addition to cancer, hDus2 may additionally be implicated in Alzheimer’s disease (28). Moreover, human Dus4L has recently been shown to be upregulated in lung adenocarcinoma and to be necessary for cell proliferation of A549 cells (29). The yeast homolog of this enzyme is known to introduce D20a and D20b; however, whether hDus4L binds tRNA and has any tRNA dihydrouridylation activity remains to be studied. Recently, dihydrouridine has been found within mRNAs and small nucleolar RNAs in budding yeast, fission yeast, and human cells, but these studies were unable to detect dihydrouridine outside of tRNA and rRNA in *E. coli* (30,31). Overall, a substantive body of research implicates dihydrouridines in tRNA to cellular fitness, health and disease.

The crystal structure of *Thermus thermophilus* Dus (*Tth*Dus; a DusA-family dihydrouridine synthase) in complex with *T. thermophilus* tRNA^Phe^ and flavin mononucleotide (FMN) has been solved (32) (Figure 1B, C). In this structure, FMN is bound in the *Tth*Dus active site surrounded by four conserved residues including K132. In the active site, FMN forms π-stacking interactions with the target nucleobase, U20, which is flipped out from the D loop into the enzyme active site. In addition to recognition by FMN, the U20 base is directly recognized by the conserved residues C93, R178, and N90. When bound to *Tth*Dus, the tRNA^Phe^ D loop is distorted at U16 and U17 in addition to U20; however, the G18-U55 and G19-C56 tertiary base pairs are maintained, suggesting the tRNA elbow is undisturbed. Indeed, no other significant tRNA conformational changes are observed within the D stem, T loop, or anticodon stem although *Tth*Dus also binds these regions (32). Thus, *Tth*Dus is likely to recognize properly folded tRNA nearing maturity. This is consistent with the lack of complex formation observed between *Tth*Dus and unmodified tRNA (32), and the fact that previous studies of purified DusA have utilized native tRNAs or bulk RNA extracted from a knockout strain rather than *in vitro* transcribed, unmodified tRNAs (12,13,18,33).

The precise reaction mechanism of tRNA dihydrouridylation by DusA remains unknown, but it is proposed that in its reduced form, the DusA-bound flavin transfers a hydride from N5 of FMNH- to C6 of the target uracil base, creating a nucleophilic centre at C5, which then attacks a conserved cysteine residue (C114 in *E. coli*, C93 in *T. thermophilus*) to obtain a proton (3,4,32). In order to recycle the flavin prosthetic group, reduced nicotinamide adenine dinucleotide (phosphate) (NADH/NADPH) binds and reduces FMNH- in an unknown manner (3,32).

Despite the widespread conservation of D20/20a and its associated dihydrouridine synthase, the molecular mechanisms, relative timing of modification, and cellular impact of *E. coli* DusA remains poorly understood. Two similarly conserved tRNA modifying enzymes, TrmA and TruB, which form 5-methyluridine (m^5^U) 54 and pseudouridine (Ψ) 55, respectively, function as tRNA chaperones by disrupting tertiary interactions between the D and T loops to access their target base, thereby providing tRNA a second chance at correctly folding (34,35). TrmA and TruB are known to act early in tRNA biogenesis where they promote tRNA modification, folding, and aminoacylation to finetune mRNA translation across several specific codons (20). In contrast to TrmA and TruB, studies of DusA homologs demonstrate a strong preference or even requirement for a modified tRNA substrate (32,36–38) and a prerequisite for proper tertiary interactions between the tRNA D and T loops for tRNA binding (32). Based on these observations we hypothesize that DusA functions in the later stages of tRNA maturation compared to the early-acting TrmA and TruB enzymes. To test this hypothesis, we characterized the molecular mechanisms of cofactor oxidation, tRNA binding, and tRNA modification using purified DusA enzyme. Additionally, we uncover the cellular functions of DusA in tRNA modification, aminoacylation, and protein translation by comparing a *dusA* deletion to its parental strain.

## Methods

### Buffers and reagents

Experiments were performed in TAKEM_4_ buffer (50 mM Tris–HCl pH 7.5, 70 mM NH_4_Cl, 30 mM KCl, 1 mM EDTA, 4 mM MgCl_2_). Oligonucleotides were purchased from Integrated DNA Technology (IDT). Unless otherwise indicated, all reagents were purchased from ThermoFisher Scientific.

### DusA overexpression and purification

The *dusA* gene was amplified from *E. coli* DH5α and inserted into the expression vector pET28a(+) using the restriction enzymes NheI and BamHI to prepare pET28a-DusA wildtype. To prepare plasmids for the expression of DusA C114A and K153A variants, site directed mutagenesis was performed using pET28a-DusA wildtype as a template with overlapping primers specified in Table 1. The coding sequences of pET28a-DusA wildtype and variants were confirmed by Sanger sequencing (Azenta). Plasmids were transformed into BL21(DE3) cells grown in LB broth supplemented with 50 µg/mL kanamycin at 37°C. Protein overexpression was induced at an OD_600_ of ∼0.6 by adding IPTG to a final concentration of 1 mM, and the temperature was reduced to 18°C. Cells were collected by centrifugation at 5000g after twenty hours, shock frozen, and stored at −80°C.

**Table 1.**
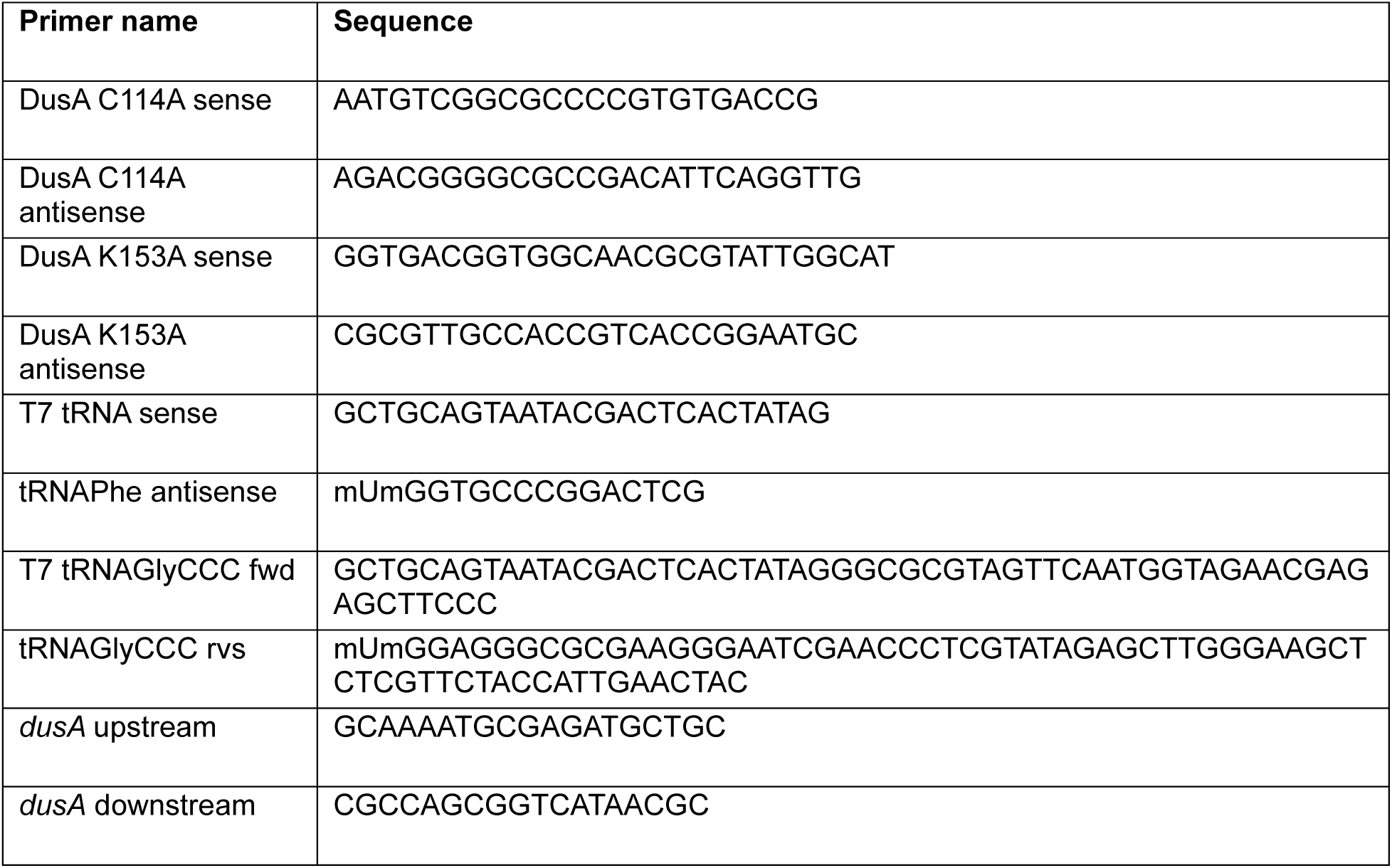
Oligonucleotide sequences used in this study.

Proteins were purified using their amino-terminal hexahistidine tag using nickel-sepharose chromatography followed by Superdex 75 size exclusion chromatography, similar to (39,40). Cells were resuspended to a final concentration in 0.2 g/mL Cell Opening Buffer (20 mM Tris-HCl pH 8.1, 400 mM KCl, 1 mM β-mercaptoethanol, 30 mM imidazole, 0.5 mM phenylmethylsulfonyl fluoride (PMSF), 5% (v/v) glycerol) and lysed for 30 minutes on ice by 1 mg/mL lysozyme. Sodium deoxycholate was added to a final concentration of 12.5 mg/g cells and incubated another 30 minutes. The solution was sonicated for four minutes total in cycles of 15 seconds on at 40% output and 45 seconds off (Fisher Scientific Sonic Dismembrator 500) and centrifuged for 30 minutes at 30,000g. The clarified lysate was loaded onto 3 mL of nickel sepharose resin (Cytiva), washed with 300 mL of Cell Opening Buffer, and DusA was eluted from the resin with Elution Buffer (same as Cell Opening Buffer except containing 500 mM imidazole and no PMSF). Following protein concentration with Amicon-15 Centrifugal Filters (Millipore), DusA was rebuffered and further purified by Superdex 75 chromatography (XK 26/100 column, Cytiva) into Superdex Buffer (20 mM HEPES-KOH pH 7.5, 150 mM KCl, 1mM β-mercaptoethanol, 0.5 mM EDTA, 5 mM MgCl_2_, 20% (v/v) glycerol). Peak fractions were concentrated, aliquoted, flash frozen, and stored at −80°C. The concentration of purified DusA was determined photometrically at A_280_ using an extinction coefficient of 41370 M^-1^ cm^-1^ (calculated with Expasy ProtParam (41)) and confirmed using comparative SDS PAGE.

Additional tRNA modifying enzymes (TrmA, TruB, IscS, and ThiI) used to prepare partially modified tRNAs were overexpressed and purified similar as above and previously published (20,35,39).

### tRNA *in vitro* transcription and modification

The DNA template for tRNA^Phe^ was prepared by PCR of plasmid pCF0 (42) using primers specified in Table 1. The DNA template for tRNA^Gly^_CCC_ was obtained by extending two overlapping primers containing the sequence of the T7 promoter and tRNA^Gly^_CCC_ by PCR (Table 1). Subsequently, *in vitro* transcription was carried out using the PCR template (10% (v/v)) in transcription buffer (40 mM Tris-HCl pH 7.5, 15 mM MgCl_2_, 2 mM spermidine, 10 mM NaCl, 10 mM DTT) with 3 mM NTPs (ATP, CTP, GTP, and UTP; Sigma), 5 mM GMP, 0.01 U/mL inorganic pyrophosphatase (Sigma), 0.3 µM T7 RNA Polymerase (purified in-house), and 0.12 U/mL RiboLock RNase inhibitor at 37°C for four hours. DNA template was degraded by addition of 2U/mL DNase I for 2 hours at 37°C. tRNA was purified by phenol/chloroform extraction to remove enzymes followed by Superdex 75 (XK 26/100) chromatography with TAKEM_4_ buffer to remove unincorporated NTPs. tRNA concentrations were determined photometrically using an extinction coefficient of 500,000 M^-1^ cm^-1^ for tRNA^Phe^ (43) and 713,400 M^-1^ cm^-1^ for tRNA^Gly^_CCC_ (IDT Oligoanalyzer).

To prepare tRNA^Phe^ and tRNA^Gly^ containing s^4^U8, m^5^U54, and Ψ55 modifications, purified tRNA (6 µM) was incubated in TAKEM_4_ buffer with 1 µM each of TruB, TrmA, IscS, and ThiI enzymes, 50 µM S-adenosylmethionine (New England Biolabs), 0.5 mM L-cysteine, 250 µM ATP, 40 µM pyridoxal 5ʹ-phosphate (Sigma), 1 mM dithiothreitol (DTT), and 0.01 U/µL RNase Inhibitor at 37°C for three hours within a total reaction volume of 7 mL. To purify partially modified tRNA, enzymes were first removed by phenol/chloroform extraction and small molecules were separated from tRNA using Superdex 75 (10/300 GL) size exclusion chromatography (44).

### DusA NAD(P)H oxidation assays to determine DusA activity and cofactor preference

To determine the NAD(P)H oxidation activity of DusA wildtype and variants, 2 µM DusA with 2 µM FMN was incubated with 500 µM NADH or NADPH in TAKEM_4_ buffer in the presence or absence of 50 µM *in vitro* transcribed tRNA^Phe^. Reactions were started by addition of NAD(P)H, and oxidation to NAD(P)+ was followed by monitoring the decrease in absorbance at 340 nm (ε_NAD(P)H_ = 6220 mM^-1^cm^-1^). Since reactions took place in aerobic conditions, oxygen acts as the final electron acceptor. To determine initial velocities (*v*_0_), the NADP+ formation was plotted versus time, and the slope of the linear portion of each reaction was determined.

### DusA tRNA dihydrouridylation assays

*In vitro* transcribed tRNA^Phe^ (5 µM) was incubated with 5 µM DusA wildtype or variant in the presence of 5 mM NADPH and 1 mM FMN in TAKEM_4_ buffer containing 1 mM DTT and 0.04 U/µL RiboLock for one hour at 37°C. Reactions were stopped by phenol extraction and modified tRNA was purified by Superdex 75 size exclusion chromatography as described above.

### Assessing DusA activity with liquid chromatography coupled to tandem mass spectrometry (LC-MS/MS)

We conducted LC-MS/MS assays to quantitatively assess the ability of our DusA protein to modify tRNA using a previously described method (45). The levels of all four standard nucleosides (A, U, G, C) and dihydrouridine (D) were measured in *in vitro* transcribed tRNA^Phe^ incubated with either no DusA, wild-type DusA, or a DusA mutant (K153A, C114). Each treated tRNA^Phe^ was hydrolyzed to mononucleotides with 300 U/μg Nuclease P1 (NEB, 100,000 U/mL), 100 mM ammonium acetate, and 100 μM zinc sulfate at 37°C overnight. The resulting mononucleotides were then dephosphorylated with 50 U/μg bacterial alkaline phosphatase (BAP, Invitrogen, 150 U/μL), 100 mM ammonium bicarbonate, and 50 mM zinc sulfate at 37°C for 5 hours. The samples were then lyophilized and resuspended in 9 μL of water and combined with 1 μL of 400 nM ^15^N_4_-inosine internal standard. Samples were separated on a Waters Acquity UPLC HSS T3 column (100 Å, 1.8 μm, 1.0 x 100 mm) heated to 35°C on an Agilent 1290 Infinity II liquid chromatography system coupled to an Agilent Technologies 6460 Triple Quadrupole mass spectrometer and ionized via electrospray ionization. Mobile phase A was 0.01% (v/v) formic acid in water and mobile phase B was 0.01% (v/v) formic acid in acetonitrile. Sample injection volume was 5 μL. Samples were run in positive mode with 4000 kV capillary voltage. The gas temperature was 350°C, the gas flow was 10 L/min, the nebulizer gas pressure was 25 psi, the sheath gas temperature was 350°C, and the sheath gas flow was 11 L/min. Calibration curves were prepared of the four canonical bases and modified nucleosides which were used to quantify sample nucleoside levels. Samples were prepared in triplicate and averaged.

### Nitrocellulose filter binding to determine affinity of DusA for tRNA

*In vitro* transcribed tRNAs were dephosphorylated with calf intestinal phosphatase (CIP; New England Biolabs) and radiolabeled using [γ-^32^P]ATP (Revvity) and T4 polynucleotide kinase (PNK; New England Biolabs). To remove unincorporated nucleotide, radiolabelled tRNA was purified through 200 µL Sephadex G-25 (Cytiva) resin in spin columns. Specific activity was determined by scintillation counting (Revvity Tri-Carb 4910TR).

Prior to the binding reaction, tRNA was refolded in 1x TAKEM_4_ buffer by heating to 65°C for five minutes and cooling to room temperature for ten minutes. Refolded tRNA (160 nM) was incubated with increasing concentrations of DusA wildtype or variant (0-50 µM) in TAKEM_4_ buffer. NADPH (1 mM) or FMN (250 µM) were added to the reactions as indicated. After incubation for ten minutes at room temperature, tRNA-enzyme mixtures were filtered through a nitrocellulose membrane under vacuum, and percent bound tRNA was determined by scintillation counting. Dissociation constants (*K*_D_) were determined by plotting percent tRNA bound (*Bound)* as a function of enzyme concentration (*[enzyme]*) and fit with a hyperbolic equation using GraphPad Prism (version 10.4):

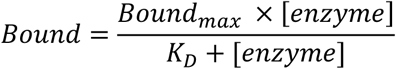

### Determination of DusA tRNA binding kinetics using stopped-flow

Fluorescein-labelled s^4^U8 tRNA^Phe^ was prepared as described previously (46,47) by first introducing the s^4^U8 modification into *in vitro* transcribed tRNA^Phe^ with ThiI and IscS enzymes, removing reaction components from s^4^U8-tRNA^Phe^ using phenol extraction followed by Superdex 75 (10/300 GL) size exclusion chromatography, and subsequently labeling the tRNA thiol group with 5-iodoacetamidofluorescein (5-IAF; Sigma). Herein, s^4^U8-tRNA^Phe^ (60 µM) was incubated with 3.2 mM 5-IAF in 12 mM HEPES-KOH pH 8.2 containing 80% (v/v) dimethyl sulfoxide at 65°C in the dark for 4 hours. To remove unincorporated fluorescent dye, successive phenol extractions were performed until the organic layer no longer appeared yellow (at least eight extractions). Trace phenol was removed by two chloroform extractions followed by ethanol precipitation, and the final fluorescent tRNA product was dissolved in water. tRNA concentration and fluorescein labeling efficiency was determined using spectrophotometry at 260 and 492 nm.

For stopped-flow experiments, 1 µM fluorescein-s^4^U8-tRNA^Phe^ was rapidly mixed with excess DusA wt or variant enzymes at indicated final concentrations (3 – 20 µM) in a KinTek SF-2004 stopped-flow at 20°C. Fluorescein was excited at 480 nm and emission was monitored from 505 nm onwards. Relative fluorescence (*Y*) was plotted against time (*t*) and fit to a two-exponential function to determine apparent rates (*k*_app_) using TableCurve 2D (version 5.01):

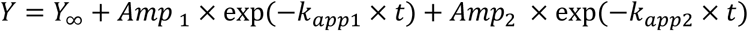

Data shown are averages of at least eight independent replicates. For DusA wt, apparent rates (*k*_app_) were plotted against the enzyme concentration ([*enzyme*]) and fit with a linear equation to determine the association rate constant, *k*_1_:

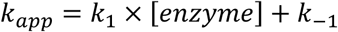

### Multiplex small RNA sequencing (MSR-seq) to determine tRNA abundance, charging, and modification

Bulk RNA was extracted using TRIzol (Invitrogen) under acidic conditions (pH <5) to preserve the tRNA aminoacyl bond from early-log phase *E. coli* BW25113 wildtype and *ΔdusA* cells (48) grown in LB medium. Strain identity was confirmed by Sanger sequencing of the *dusA* locus using primers specified in Table 1. After shock freezing wet cell pellets from 15 mL culture grown to ∼0.4 OD_600_, cells were opened with 3 mL TRIzol and 0.6 mL chloroform was added after a three-minute incubation, according to the manufacturer’s protocol. After centrifugation (5000g, 45 minutes), the aqueous phase was transferred to a new tube and ethanol precipitated several times. Pure RNA was resuspended in 10 mM NaOAc (pH 4.8) and stored at −80 °C. Concentrations were determined using UV spectrometry (NanoDrop 2000c). Three biological replicates from each strain were isolated.

Library preparation, MSR-tRNA-seq, and bioinformatics were performed by MesoRNA (Chicago, IL, USA). Reads were demultiplexed and trimmed as appropriate in accordance with the MSR-seq library preparation protocol (49). Quality control was performed with FastQC (https://www.bioinformatics.babraham.ac.uk/projects/fastqc/). Reads were aligned to tRNA sequences adapted from GtRNAdb (50) with bowtie2 (51). Further analysis and plots were generated with custom software developed by MesoRNA.

### Tandem codon GFP reporter assays

BW25113 wildtype and *ΔdusA* strains were transformed with reporter plasmids carrying an arabinose-inducible fluorescent transcriptional cassette encoding superfolder green fluorescent protein (sfGFP) followed by mCherry fluorescent protein (kind gift from Assaf Katz, Universidad de Chile, Santiago, Chile) (52). Each reporter contains a set of four tandem repeats of a codon following the third sfGFP codon, such that sfGFP expression relies on translation of the four repeated codons. To account for subtle differences in translation between strains, plasmid S1 was used as a control, which lacks any additional codons in sfGFP. sfGFP and mCherry fluorescence were determined similarly as previously described (20,52). In brief, overnight cultures were diluted 1:20 to an OD_600_ of ∼0.1 and grown in LB medium supplemented with 100 µg/mL ampicillin until reaching an OD_600_ between 0.4-0.6. Cells were then diluted 1:4 into fresh LB medium containing 100 µg/mL ampicillin and 0.4% (w/v) arabinose in black, optical bottom 96 well plates and grown at 37°C with shaking at 100 rpm. Three-hours post induction, OD_600_, sfGFP fluorescence (excitation: 470-15 nm, emission: 515-20 nm), and mCherry fluorescence (excitation: 570-15 nm, emission: 620-20 nm) were measured using a CLARIOstar Plus plate reader (BMG Labtech). sfGFP/mCherry ratios for each test codon were normalized to the sfGFP/mCherry ratio of plasmid S1 for the respective strain:

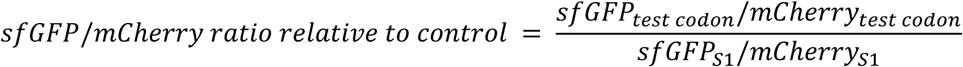

For each codon, translation between strains was compared using two-way ANOVA. Significant differences (*p* < 0.05) between strains are indicated.

## Results

Although dihydrouridine is one of the most frequent tRNA modifications, the enzymes that form dihydrouridine remain understudied in comparison to other tRNA modifying enzymes. Here, we examined the molecular mechanism of *E. coli* DusA and clarify the roles of this enzyme and D20/D20a modifications *in vivo* during tRNA maturation and protein synthesis.

### DusA can form dihydrouridine in *in vitro* transcribed tRNA^Phe^

Previous work has shown that some tRNA dihydrouridine synthases require the presence of previously introduced tRNA modifications for tRNA modification and/or NAD(P)H reduction activity (32,36), and previous studies of DusA have utilized native tRNAs rather than unmodified transcripts (12,13,18,33). Asking whether *E. coli* DusA also required a partially modified tRNA substrate for activity, we incubated a high concentration (5 µM) of purified DusA wildtype or variant with a stoichiometric amount of unmodified tRNA for one hour, purified the resulting DusA-modified tRNA and determined the amount of dihydrouridine formed by mass spectrometry. We found that DusA wildtype can modify *in vitro* transcribed tRNA^Phe^ under the conditions tested, with an end level of 5.65% D/total U content (Figure 1D). DusA has previously shown to be specific to U20 in tRNA^Phe^ (13,18), which has 16 U nucleotides in *E. coli* tRNA^Phe^, i.e. 5.65% D/total U accounts for >90% of U20 being converted to D20. Similar to previous reports, substitution of the proposed catalytic cysteine residue to alanine (DusA C114A) resulted in a drastic decrease in dihydrouridylation by DusA (Figure 1D). Previous work has shown that the DusA K153A variant or its equivalent in other organisms cannot form D20 when expressed in cells (33,53). In contrast to these *in vivo* experiments, in our *in vitro* assay, DusA K153A was able to form dihydrouridine in *in vitro* transcribed tRNA, although with a lower end-level than that of the wildtype enzyme, suggesting that dihydrouridine formation is slower for this variant compared to the wildtype enzyme. Based on previous studies of the equivalent variant of *Tth*Dus, we hypothesize that catalysis by DusA K153A is slow due to decreased affinity for FMN and tRNA (32). Indeed, whereas DusA C114A co-purifies with a flavin cofactor similar to the wildtype enzyme, no absorbance at ∼375 nm and ∼450 nm, characteristic of a flavoprotein was observed for purified DusA K153A, suggesting impaired FMN interaction for this variant (Figure 1E).

### Characterization of NAD(P)H oxidation by DusA

Whereas most characterized prokaryotic and eukaryotic tRNA dihydrouridine synthases oxidize NADPH faster than NADH (15,16,37), the preferred FMN reduction substrate for DusA has not yet been reported. To address this, we monitored the oxidation of NAD(P)H by measuring the loss of absorbance at 340 nm as NAD(P)H is oxidized to NAD(P)+. Under multiple turnover conditions with 500 µM NAD(P)H, DusA quickly oxidizes NADPH in the absence and presence of tRNA, with initial velocities of 85 ± 3 µM min^-1^ and 68 ± 4 µM min^-1^, respectively (Figure 1F, Table 2). Thus, DusA can oxidize NADPH, with air acting as the final electron acceptor, even in the absence of tRNA. In contrast, the oxidation of NADH is significantly slower, but still above the background (Figure 1F, Table 2).

Next, we asked if the DusA C114A and DusA K153A variants can oxidize NADPH although DusA C114A was not active in tRNA modification and DusA K153A purified without the FMN cofactor (Figure 1G). DusA C114A retained the ability to oxidize NADPH, albeit with a 5-fold slower initial velocity compared to DusA wildtype (Table 2). NADPH oxidation by DusA K153A was even slower than that for DusA C114A, but still above background with a 25-fold slower initial velocity compared to the wildtype enzyme, suggesting some residual NADPH reduction activity remains for this variant (Table 2).

**Table 2.** Initial velocity (*v*_0_) of NAD(P)H oxidation by DusA wildtype and variants.

| | NADPH oxidation $v_0$, μM min<sup>-1</sup> | NADH oxidation $v_0$, μM min<sup>-1</sup> |
| --- | --- | --- |
| **DusA wt** | 85 ± 3 | 3.8 ± 0.7 |
| **DusA wt + tRNA<sup>Phe</sup>** | 68 ± 4 | ND |
| **DusA C114A** | 17 ± 0.5 | ND |
| **DusA K153A** | 3.5 ± 0.2 | ND |

### Characterization of tRNA binding by DusA

Next, we investigated the affinity of DusA for two of its known substrate tRNAs: tRNA^Phe^ and tRNA^Gly^_CCC_. Previous work with *Tth*Dus has suggested this enzyme requires its substrate tRNA to contain modifications for efficient binding and/or catalysis, at least at high temperatures (32,38), whereas yeast Dus2 requires tRNA to be modified for efficient NADPH oxidation activity (36), and human Dus2 does not require tRNA to be modified for binding, but does require previous modifications for enzymatic activity (37). Since all previous studies of *E. coli* DusA utilized tRNA substrates that were isolated from a knockout strain (13,18,33), we sought to clarify whether modifications are important for the binding of DusA to tRNA. For this reason, we also prepared partially modified tRNA^Phe^ and tRNA^Gly^_CCC_ that contain s^4^U8, m^5^U54, and Ψ55 by modifying *in vitro* transcribed tRNAs with ThiI, IscS, TrmA, and TruB (44). For tRNA^Gly^_CCC_, these constitute all of its known native modifications except for the DusA-introduced D20 modification, and for tRNA^Phe^ these are three of its nine total non-D20 modifications (54). The affinity of wildtype DusA for unmodified tRNA^Phe^ was found to be 7 ± 1 µM, whereas the affinity of DusA for unmodified tRNA^Gly^_CCC_ was about two-fold lower at 14 ± 2 µM (Figure 2A, B, Table 3). Presence of modifications within tRNAs slightly but significantly increase the affinity of DusA for tRNA^Phe^ and tRNA^Gly^_CCC_ with dissociation constants of 5.9 ± 0.3 µM and 9.3 ± 0.7 µM, respectively (Figure 2A, B, Table 3). We next assessed whether cofactor presence may stabilize tRNA binding, thereby increasing the affinity. DusA co-purifies with FMN, but it is unknown whether this cofactor is present at stoichiometric amounts. Thus, we examined DusA binding to tRNA^Phe^ and tRNA^Gly^_CCC_ in the presence of excess FMN or NADPH. Surprisingly, addition of either cofactor significantly reduced tRNA binding with addition of NADPH lowering the affinity up to ∼1.5-fold and FMN lowering the affinity more than 5-fold (Figure 2A, B, Table 3). Corresponding with this increase in *K*_D_, the end level of tRNA binding is lower upon cofactor addition compared to the absence of cofactors.

**Table 3.** Dissociation constants (*K*_D_) for DusA wt and variants binding to tRNA^Phe^ and tRNA^Gly^ in the presence or absence of enzyme cofactors.

| | tRNA<sup>Phe</sup> $K_D$, $\mu\text{M}$ | tRNA<sup>Gly</sup><sub>CCC</sub> $K_D$, $\mu\text{M}$ |
| --- | --- | --- |
| **Unmodified tRNA** | $7.3 \pm 0.8 \mu\text{M}$ | $13.8 \pm 2.3 \mu\text{M}$ |
| **Modified tRNA** | $5.9 \pm 0.3 \mu\text{M}$ | $9.3 \pm 0.7 \mu\text{M}$ |
| **Unmodified tRNA + NADPH** | $10.5 \pm 0.9 \mu\text{M}$ | $9.4 \pm 0.9 \mu\text{M}$ |
| **Unmodified tRNA + FMN** | $>50 \mu\text{M}$ | $>50 \mu\text{M}$ |
| **DusA C114A** | $17.6 \pm 1.1 \mu\text{M}$ | ND |
| **DusA K153A** | $>50 \mu\text{M}$ | ND |

**Figure 2.**
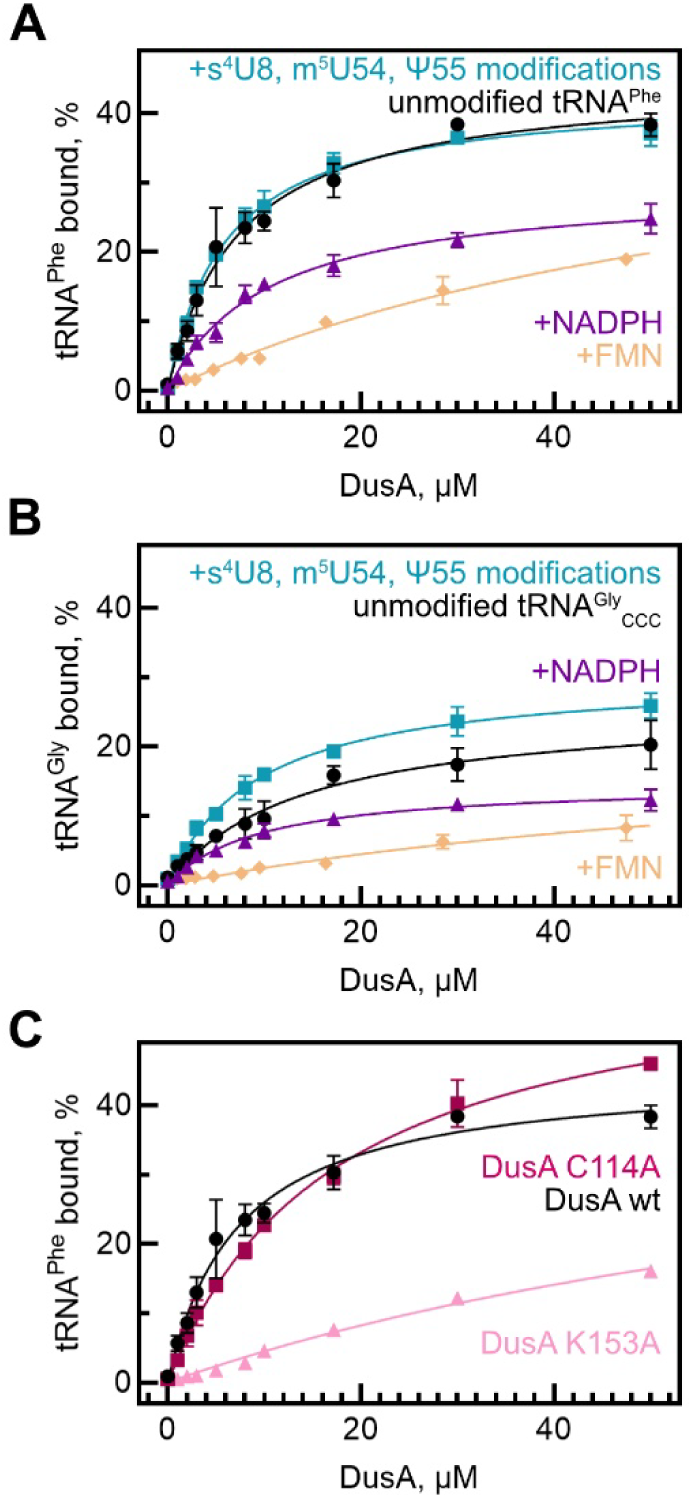
Affinity of DusA for tRNA. **(A)** Nitrocellulose filtration titration for DusA wt binding to unmodified tRNA^Phe^ (black circles) or to partially modified tRNA^Phe^ (teal squares) in the absence of added cofactors. Nitrocellulose filtration data for DusA wt binding to unmodified tRNA^Phe^ in the presence of 1 mM NADPH (purple triangles) or 40 µM FMN (cream diamonds). **(B)** Nitrocellulose filter binding data for DusA wt binding to unmodified or partially modified tRNA^Gly^_CCC_, similar to panel A. **(C)** Nitrocellulose filtration for DusA C114A (dark pink squares) and K153A (light pink circles) binding to unmodified tRNA^Phe^. For comparison, the binding curve for wildtype DusA shown in panel A is repeated here (black circles). N=3 for each curve and error bars represent standard deviations. Dissociation constants for each data set in panels A-C were determined by hyperbolic fitting and are displayed in Table 3.

Next, we examined whether the DusA variants are deficient in tRNA binding. DusA C114A binds to tRNA^Phe^ with a *K*_D_ around 18 µM (∼2-fold weaker than DusA wildtype) with approximately the same end level of binding as the wildtype enzyme. In contrast, DusA K153A was found to bind tRNA^Phe^ with a much lower affinity, with a *K*_D_ above 50 µM and observed end level approximately half of that for the wildtype enzyme (Figure 2C, Table 3).

### Rapid kinetic dissection of DusA binding to tRNA

In order to monitor the binding of DusA to tRNA^Phe^ in real time, we utilized a previously established fluorescent tRNA stopped flow assay (46,47). Herein, we introduced the s^4^U8 modification within *in vitro* transcribed tRNA^Phe^ and subsequently labeled the thiol group with fluorescein (Figure 3A). Rapid mixing of DusA wildtype (20 µM) with fluorescent tRNA (1 µM) results in a two-phase decrease fluorescence within one second (black trace, Figure 3B). Fitting this data with a two-exponential function reveals a fast phase with an apparent rate of 146 ± 17 s^-1^ and a slow phase with an apparent rate 17 ± 2 s^-1^, respectively (Figure 3B). Similar to wildtype DusA, rapid mixing of 20 µM DusA C114A and K153A variants with fluorescent tRNA also results in a two-phase fluorescence decrease (Figure 3B, C); however, the amplitude of these fluorescence decreases are significantly smaller than that for the wildtype enzyme. Fitting with a two-exponential function yields apparent rates equal to 91 ± 11 s^-1^ and 15 ± 1 s^-1^ for DusA C114A and 29 ± 2 s^-1^ and 1 ± 3 s^-1^ for DusA K153A (Figure 3B, 3C).

**Figure 3.**
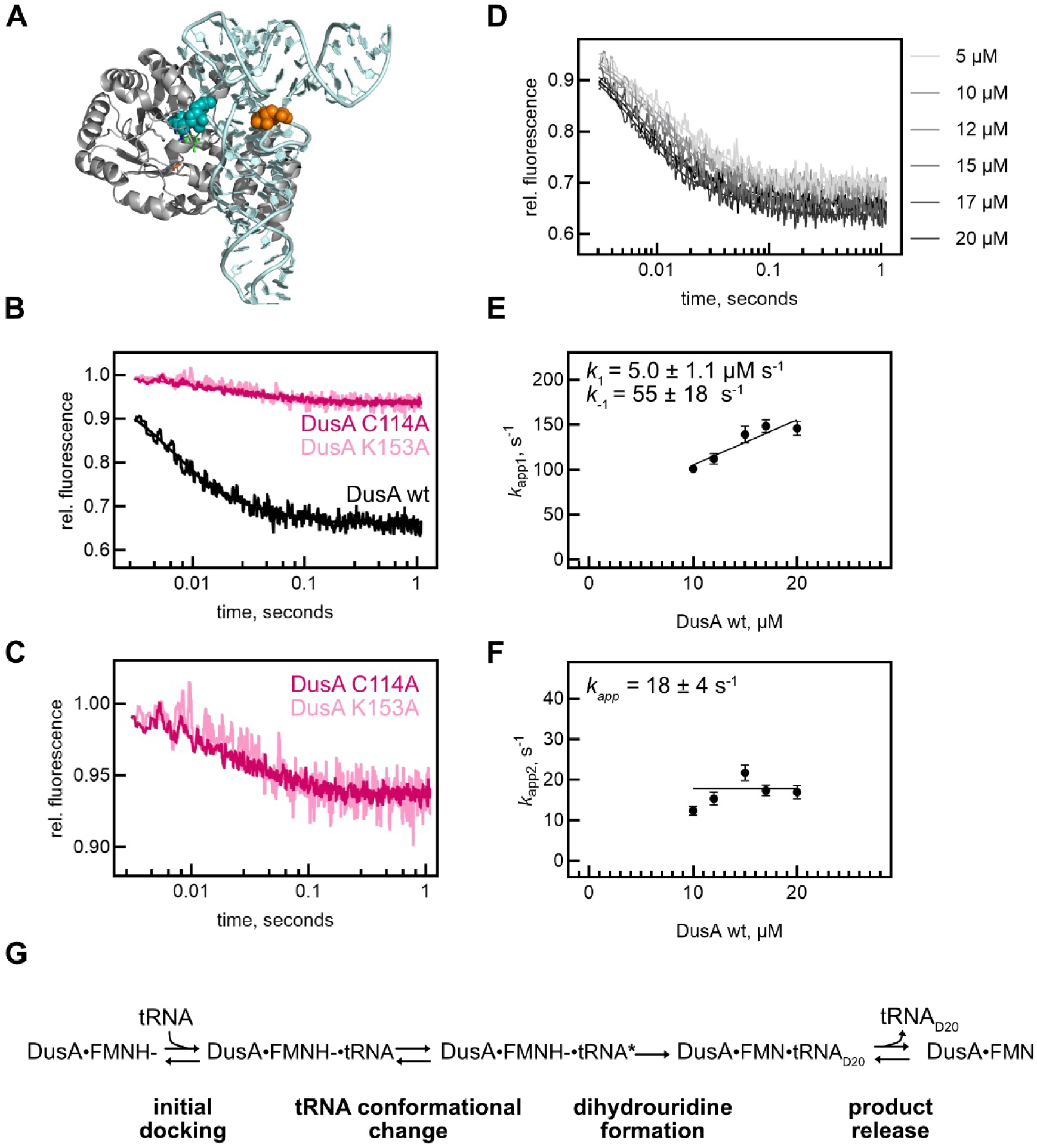
Rapid kinetic dissection of DusA binding to tRNA^Phe^. **(A)** Structure of *T. thermophilus* Dus (grey) bound to tRNA^Phe^ (light teal) (PDB 3B0V) (32) showing the locations of the target nucleotide of DusA (U20) as teal spheres and of the fluorescently labeled nucleotide (s^4^U8) as orange spheres. **(B)** Rapid mixing of 20 µM DusA wt (black curve), DusA C114A (dark pink curve), or K153A (light pink curve) with 1 µM fluorescent tRNA^Phe^ at 20°C. Curves were fit to two-exponential equations. **(C)** Zoomed-in view of rapid binding of DusA C114A and K153A variants to fluorescently labeled tRNA^Phe^ to better visualize the small decrease in fluorescence. **(D)** Pre-steady state binding of increasing DusA wt concentrations to 1 µM fluorescent tRNA^Phe^ to determine apparent rates. Plotting *k*_app_ against time shows that *k_app1_* is concentration dependent **(E)**, whereas *k*_app2_ is independent of DusA concentration **(F)**. **(G)** Kinetic mechanism for tRNA binding to DusA. Following binding of tRNA by DusA, conformational changes take place.

To gain a deeper understanding of the kinetics of wildtype DusA binding to tRNA, the stopped-flow experiments were repeated at different concentrations of DusA (Figure 3E, 3F), and apparent rates were plotted against DusA concentration (Figure 3E, 3F). We observed that the apparent rate corresponding to the fast phase (*k_app1_*) is dependent on DusA concentration, consistent with this phase representing a bimolecular binding event between DusA and tRNA. Linear fitting reveals a *k*_1_ of 5 ± 1 µM^-1^ s^-1^ and an estimated *k*_off_ equal to 55 ± 18 s^-1^ (Figure 3E), which agrees with the determined *K*_D_ of ∼7 µM (Figure 2A, Table 3). The apparent rate corresponding to the slow phase (*k_app2_*) was found to be not dependent upon the DusA concentration, with an average *k_app_* of ∼18 s^-1^ (Figure 3F). This second phase therefore reflects a unimolecular event such as a conformation change of tRNA. Taken together, this stopped-flow data suggests that DusA binds tRNA with a two-step mechanism, with the first step encompassing binding and the second step encompassing a conformational change (Figure 3G).

### Impact of DusA on tRNA abundance and charging *in vivo*

To date, there are no reported growth defects for the *E. coli ΔdusA* strain, and the conserved biological functions for DusA binding and modifying many tRNAs remains unclear. To address this question, we utilized multiplex small RNA sequencing (MSR-seq), to compare the abundance, aminoacylation, and modification status for each tRNA isoacceptor between the wildtype and *ΔdusA* strain (49). Herein, bulk RNA isolated in acidic conditions was subjected to periodate oxidation and β-elimination followed by library preparation and Illumina sequencing. Comparison of the abundances of individual tRNA species between the wildtype and *ΔdusA* strain revealed only one tRNA was significantly upregulated in *ΔdusA* (tRNA^Leu^_CAA_, Figure 4A). Although no other tRNAs were found to be individually significantly different in the knockout strain compared to wildtype, on a global scale, more tRNAs are upregulated upon *dusA* deletion compared to downregulated (Figure 4A). Indeed, on average, each tRNA is 1.3-fold more abundant in the *ΔdusA* strain than in wildtype (Figure 4B).

**Figure 4.**
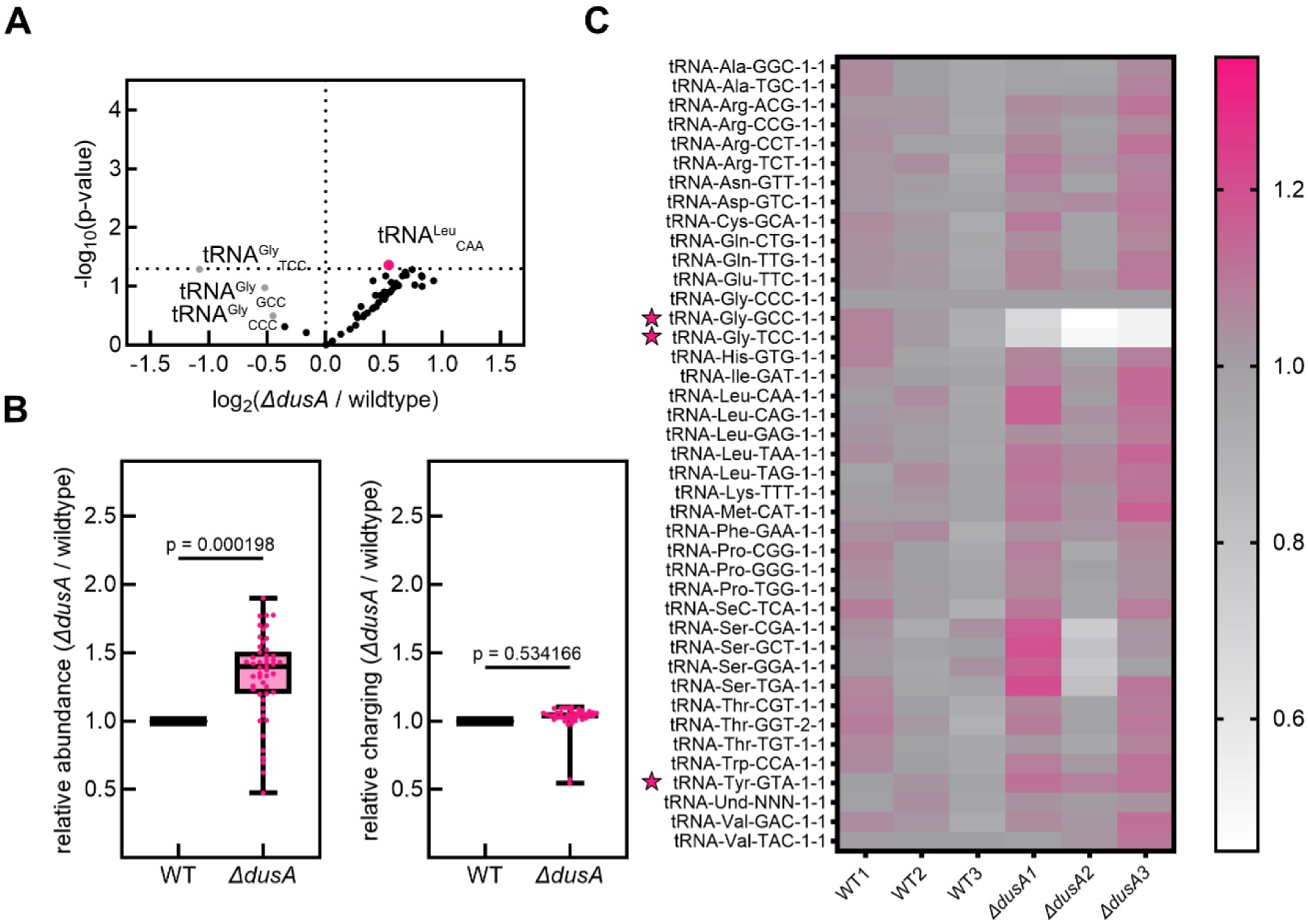
Abundance and aminoacylation levels of certain tRNAs are affected by DusA. **(A)** tRNA abundance change in *ΔdusA* compared to the wildtype strain (n = 3 for each strain). Horizontal dashed line indicates *p* = 0.05. One tRNA with significant changes in the knockout strain was identified and is indicated by a large pink circle, tRNA^Gly^ isoacceptors are indicated by grey color **(B)** Box and whisker plot showing relative abundance level changes (left) and relative charging level changes (right) for all tRNAs in wildtype and *ΔdusA* strains. **(C)** Heatmap of relative charging level for each tRNA in all replicates for wildtype and *ΔdusA* strains. Stars indicate significant (*p* < 0.05) differences in tRNA charging between the wildtype and *ΔdusA* strains.

Next, we examined the fraction of each tRNA charged between the wildtype and *ΔdusA* strains. In contrast to our previous study with *trmA* and *truB* (20), deletion of *dusA* does not globally affect tRNA aminoacylation in the *ΔdusA* strain (Figure 4B). Instead, the charging fraction of only a few tRNAs was changed as we observed aa significant decreased for tRNA^Gly^_GCC_ and tRNA^Gly^_UCC_ and a significant increase for tRNA^Tyr^_GTA_ charging within the *ΔdusA* strain compared to wildtype (Figure 4C). Intriguingly, one of the two affected tRNA^Gly^ isoacceptors tRNA^Gly^_UCC_, has not been previously identified to be modified by DusA (18,54).

To determine whether the translation of specific codons is affected by the altered levels of tRNA abundance or charging of specific tRNAs in the *ΔdusA* strain, we used a sfGFP-based codon reporter, wherein four tandem repeats of a specific codon are present near the beginning of the sfGFP open reading frame and mCherry is independently translated as a control (Figure 5A) (52). First, we examined translation of Leu codons, as the abundance of tRNA^Leu^_CAA_ is significantly higher in the knockout strain compared to wildtype. No significant difference in sfGFP expression for the reporter containing its cognate Leu codon, TTG, was observed between the two strains (Figure 5B). However, we noticed a decrease in sfGFP expression in *ΔdusA* for the reporter containing tandem repeats of the Leu codon CTA, read by DusA-modified tRNA^Leu^_TAG_, which did not display a change in abundance or charging in the knockout strain (Figure 5B).

**Figure 5.**
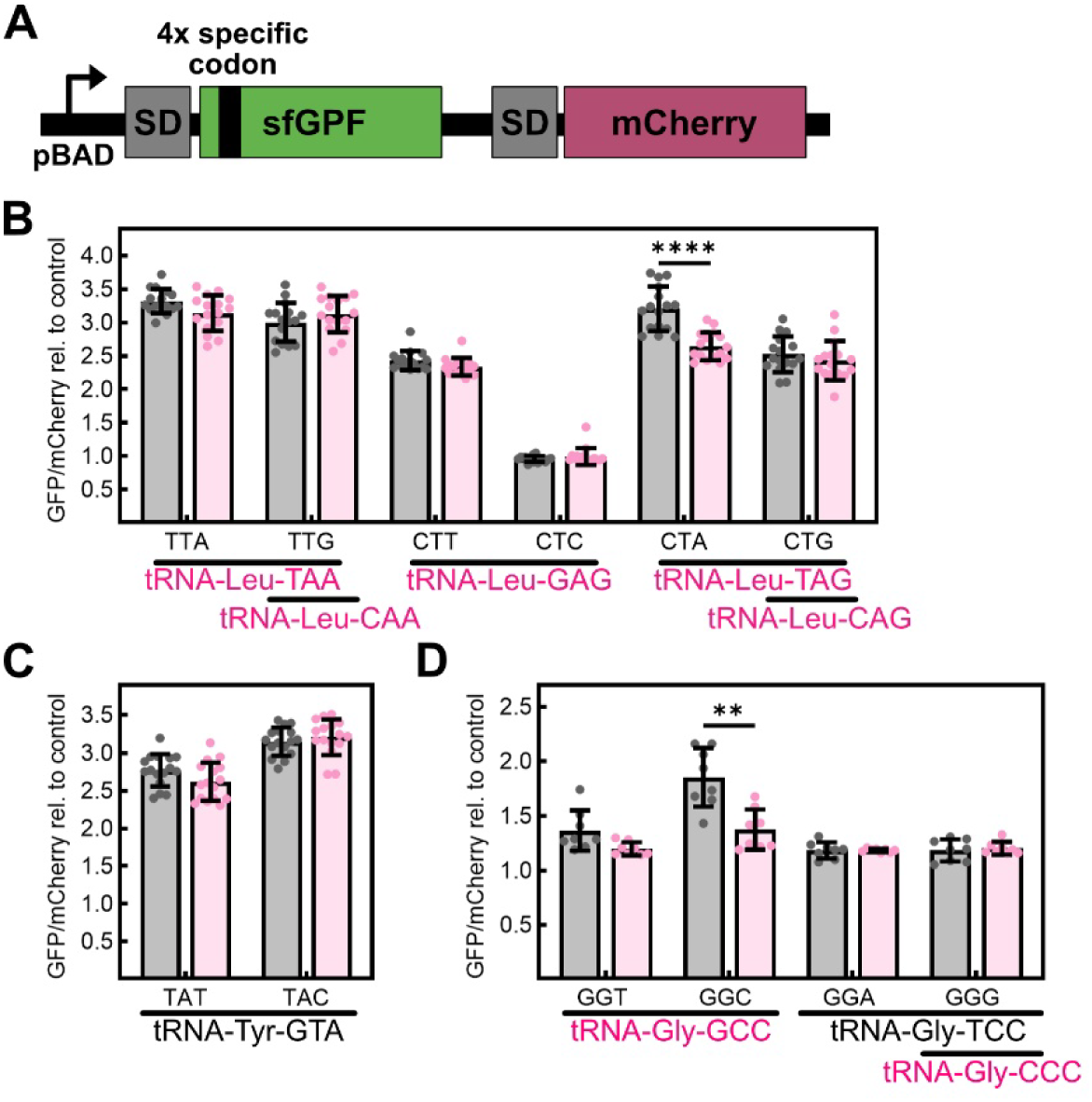
Changes in tRNA abundance and aminoacylation are not correlated with translational changes in the *ΔdusA* strain. Measurements of codon specific translation using a sfGFP codon reporter library wherein four tandem repeats of a specific codon are present near the beginning of the sfGFP open reading frame. sfGFP and mCherry expression is induced in early log phase. Whereas sfGFP expression is dependent on readthrough of the four repeated codons, mCherry is independently expressed and is used for normalization. SD: Shine Dalgarno sequence, figure adapted from (52) **(A)**. sfGFP/mCherry expression ratios for all Leu **(B)**, Tyr **(C)**, and Gly **(D)** codons. The cognate tRNA(s) are indicated underneath each codon, coloured in pink if known to be modified by DusA and black if not known to contain D20/D20a (18). For each codon, at least eight biological replicates were measured. * Indicates *p* < 0.05, ** indicates *p* < 0.01, and *** indicates *p* < 0.001.

Additionally, we examined translation of the four Gly codons, as tRNA charging was found to be reduced in the *ΔdusA* strain for two of the three tRNA^Gly^ isoacceptors. Indeed, the translation of the Gly codon GGC is significantly decreased in *ΔdusA* compared to wildtype (Figure 5C). This codon is read by tRNA^Gly^_GCC_, which is one the isoacceptors with reduced aminoacylation in the deletion strain, and which is known to be modified by DusA (18). This tRNA additionally reads Gly GGT as a cognate codon; however, the translation of Gly GGT is not altered in *ΔdusA*. Although charging tRNA^Gly^_UCC_ was also found to be decreased in the knockout strain, the translation of both of its cognate codons is not affected by loss of *dusA* (Figure 5C). Likewise, the translation of both Tyr codons is not affected by *dusA* deletion, although the charging of the cognate tRNA is slightly but significantly increased in the knockout strain (Figure 5D).

From the MSR-seq experiment, we were additionally able to determine relative changes in certain modification levels between the two strains for modifications that leave a reverse transcriptase signature (49). For example, acp^3^U47 is located nearby to D20 in the tRNA elbow (Figure 6A, 6B) and this modification results in deletions and mutations during reverse transcription that are read during sequencing (Figure 6C, D). We examined the sum of the relative fraction of tRNA reads that contain deletion and mutations at position 47 for the seven tRNAs known to contain acp^3^U47 (55,56) (Figure 6E). Of these tRNAs, six are known to be also modified by DusA (18). For each of these tRNAs except tRNA^Val^_GAC_, the sum of deletions and mutations at position 47 was lower for tRNAs extracted from the *ΔdusA* strain compared to wildtype, with tRNA^Met^_CAT_, tRNA^Ile^_CAT_, and tRNA^Arg^_ACG_ showing significant reductions (Figure 6E). This observation indicates that the absence of D20 also reduces the presence of acp^3^U47 in several tRNAs.

**Figure 6.**
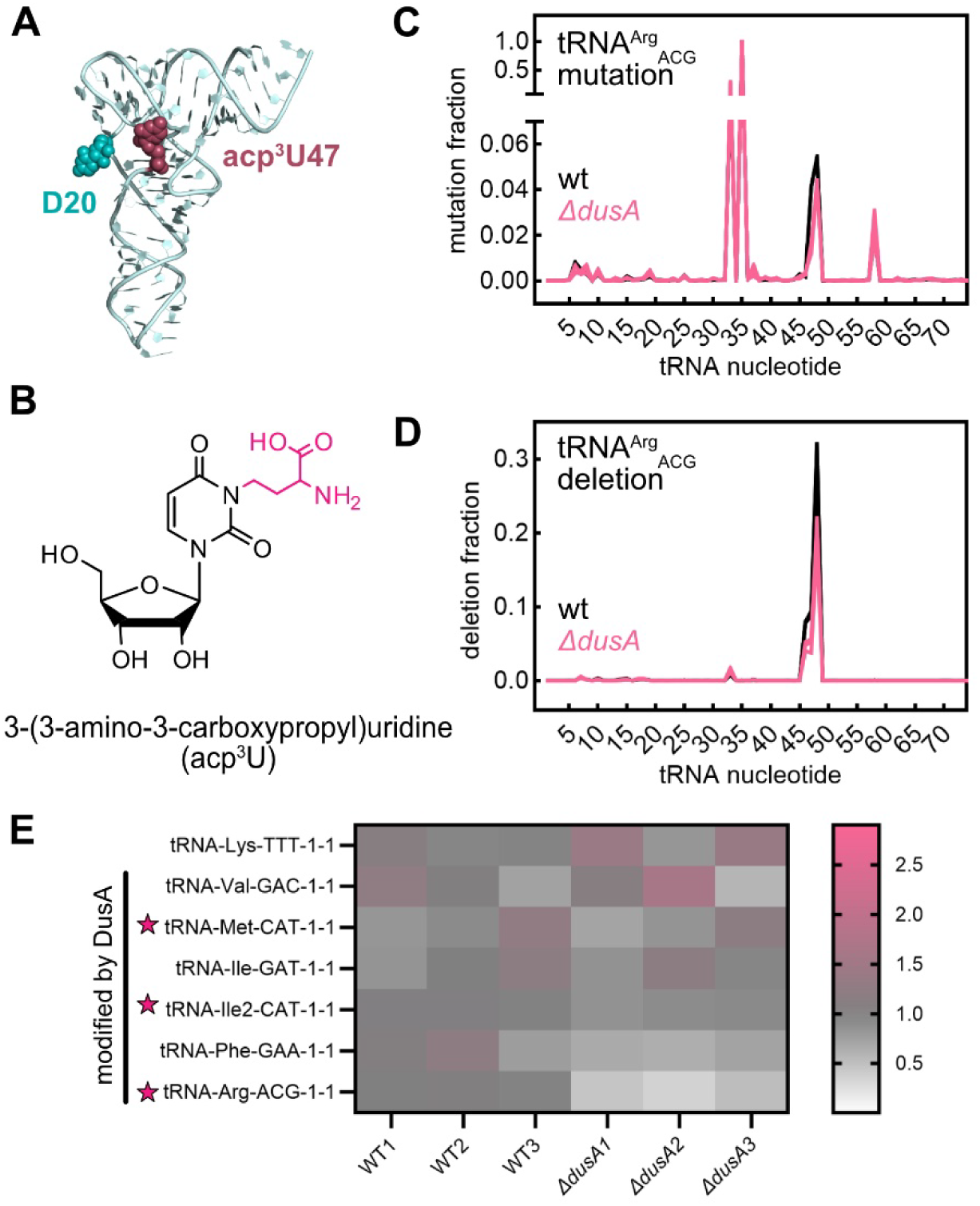
Changes in acp^3^U47 content in *ΔdusA E. coli*. **(A)** Structure of unmodified *E. coli* tRNA^Phe^ (PDB 3L0U) (57) showing relative locations for D20 (teal spheres) and acp^3^U47 (burgundy spheres). **(B)** Chemical structure of acp^3^U with the modified portion of the nucleobase highlight in pink. Relative mutation fraction **(C)** and deletion fraction **(D)** comparing wildtype (grey) and *ΔdusA* (pink) at each position in tRNA^Arg^_ACG_. **(E)** Heatmap showing the relative sum for mutation and deletion fractions at position 47 for all *E. coli* tRNAs known to contain acp^3^U47.

Moreover, we examined changed in the abundance of I34 in tRNA^Arg^_ACG_. This tRNA editing event causes reading of position 34 as a G rather than an A, resulting in an apparent mutation during tRNA mapping (Figure 6C). No change in abundance for this essential tRNA anticodon modification was observed between the wildtype and *ΔdusA* strains, with nearly 100% of this position read as a G in both strains (Figure 6C). We additionally examined for changes in mutations and deletions at positions 32 and 37 but did not identify any potential changes in tRNA modifications at these sites (data not shown).

### DusA affects the translation of certain codons

Intriguingly, when we examined specific translation for codons that had cognate tRNAs with altered abundance or aminoacylation in *ΔdusA* compared to wildtype, we found that not all of these codons display significant changes in translation, whereas certain codons read by unaffected tRNAs show differences in translation (Figure 5). Together with the fact that levels of at least acp^3^U47 are changed in other tRNAs (Figure 6), we wondered if the presence of the *dusA* gene may affect overall translation or the specific translation of additional codons. Thus, we examined codon specific translation using the sfGFP reporter as above with several additional codons (Figure 7).

**Figure 7.**
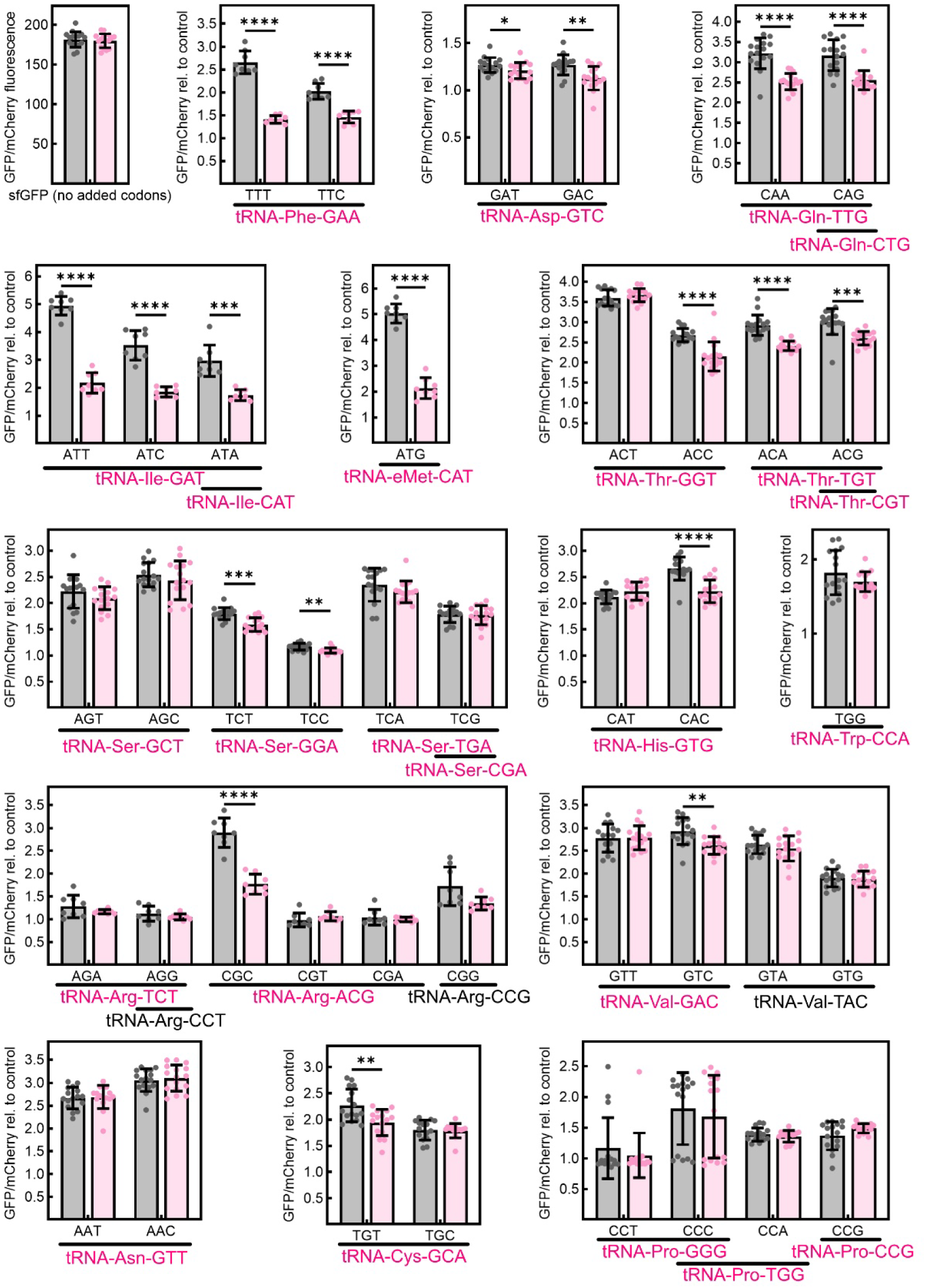
Translation of several codons read by DusA-modified tRNAs are decreased in the *ΔdusA* strain. Relative sfGFP/mCherry expression ratios for indicated codons. The cognate tRNA(s) are indicated underneath each codon, coloured in pink if known to be modified by DusA and black if not known to contain D20/D20a (18). For each codon, at least eight biological replicates were measured. * Indicates *p* < 0.05, ** indicates *p* < 0.01, and *** indicates *p* < 0.001.

To assess whether there is a difference in global translation between the wildtype and *dusA* knockout strain, we compared sfGFP expression between the two strains in the absence of additional codons as a control. No significant difference in sfGFP translation was observed (Figure 7). However, we did observe several significant codon-specific decreases in translation in the *ΔdusA* strain compared to the wildtype strain for several codons including Phe UUU and UUC (tRNA^Phe^_GAA_), Asp GAC and GAT (tRNA^Asp^_GTC_), Met ATG (tRNA^eMet^_CAT_), Ser TCT and TCC (tRNA^Ser^_GGA_), Ile ATT, ATC, and ATA (tRNA^Ile^_GAT_ and/or tRNA^Ile^_GAT_), Gln CAA and CAG (tRNA^Gln^_TTG_ and/or tRNA^Gln^_CTG_), and Thr ACA and ACG (tRNA^Thr^_TGT_, and/or tRNA^Thr^_CGT_) codons (Figure 7). Notably, for these tRNAs, the translation of each of their cognate codons is decreased when *dusA* is deleted. For other tRNAs, including tRNA^Val^_GAC_, tRNA^His^_GTG_, tRNA^Cys^_GCA_, tRNA^Arg^_ACG_, and tRNA^Thr^_GGT_, translation of only one of the two or three cognate codons is affected by the loss of *dusA* (Figure 7). In other cases, none of the cognate codons read by tRNAs known to be modified by DusA are affected by *dusA* knockout, such as for tRNA^Asn^_GTT_, tRNA^Trp^_CCA_, tRNA^Ser^_GTC_, tRNA^Ser^_TGA_, tRNA^Ser^_CGA_, tRNA^Arg^_TCT_, and all tRNA^Pro^ isoacceptors (Figure 7). Importantly, all these listed tRNAs are known to be modified by DusA (18). We additionally tested several codons read by tRNAs that are not modified by DusA including tRNA^Val^_TAC_, tRNA^Arg^_CCT_, tRNA^Arg^_CCG_, and tRNA^Tyr^_GTA_. As expected, knockout of *dusA* does not affect the translation of any of these cognate codons (Figure 5, Figure 7). Taken together, our results suggest that DusA positively impacts the translation of many, but not all, cognate codons for DusA-modified tRNAs.

## Discussion

Herein, we uncover novel insights into the mechanisms and cellular functions of *E. coli* DusA. These results illuminate key differences between DusA and two other highly conserved tRNA modifying enzymes, TrmA and TruB. In summary, our experiments demonstrate for the first time that *E. coli* DusA can bind and modify unmodified tRNA, though it binds more tightly to tRNA that already contains modifications. tRNA binding by DusA occurs via a two-step mechanism, with the second step constituting a conformational rearrangement in tRNA. Finally, we demonstrate that DusA has distinct cellular functions from TrmA and TruB, wherein DusA acts in the intermediate or late stages of tRNA maturation on a mostly mature and properly folded tRNA substrate to directly improve the function of specific tRNAs during mRNA translation on the ribosome without broadly affecting tRNA abundance or aminoacylation.

### Molecular determinants of DusA substrate binding and activity

To clarify the molecular determinants of DusA, we examined the roles of two residues, C114 and K153, previously found to be necessary for overall dihydrouridine formation (32,33). Despite being strongly impaired in tRNA modification, DusA C114A remains able to oxidize NADPH and bind tRNA albeit with reduced activity and affinity, respectively. In contrast, we found that DusA K153A can still generate dihydrouridine in tRNA *in vitro* even though it is drastically impaired in NADPH oxidation and tRNA binding and does not co-purify with FMN in agreement with its low affinity for the flavin cofactor (32). However, in the cellular context, this variant is unable to form detectable amounts of dihydrouridine (33,53). Taken together, these findings validate distinct roles for C114 and K153, with C114 necessary for catalysis, likely acting as a general acid as previously proposed, whereas K153 is required for FMN binding and thereby stable tRNA binding (32).

Additionally, for the first time we clarify that NADPH is preferred source of reducing equivalent for FMN regeneration by DusA (Figure 1F). This finding is in line with experiments with several dihydrouridine synthases, with only one enzyme identified so far to prefer NADH (*Bacillus subtilis* DusB2) (15,16,37)

### tRNA modifications contribute to DusA binding and activity during the intermediate stages of tRNA maturation

Although all previous studies of *E. coli* DusA have utilized native tRNAs containing all modifications except for D20 (12,13,18,33), we demonstrate here that DusA can bind and modify *in vitro* transcribed tRNA^Phe^ (Figure 1D, Figure 2). Notably, we find that DusA consistently binds tRNA^Phe^ more than 1.5-fold tighter that tRNA^Gly^_CCC_ regardless of its modification status.

This suggests that DusA engages different tRNA species with distinct affinities, perhaps reflecting differences in primary sequence or intrinsic structural stability of the elbow region, which DusA homologs are known to recognize (32,58).

For both tRNAs examined, we find that presence of s^4^U8, m^5^U54, and Ψ55 modestly but significantly enhance the affinity of DusA. The binding preference of DusA for a modified tRNA substrate contrasts TrmB, which is not sensitive to tRNA modification status, and TruB, which prefers binding unmodified tRNA over modified forms (40). TrmA behaves similarly to DusA in terms of binding, preferring to bind tRNA with Ψ55 over unmodified tRNA; however TrmA displays slower steady-state methylation kinetics for Ψ55 tRNA (40). Because we measured the end-level of dihydrouridine formation in unmodified tRNA, it remains to be determined whether the presence of previous modifications affects how fast DusA modifies tRNA.

Based on our observations and previous studies of DusA homologs, we speculate that *E. coli* DusA will modify tRNA that already contains modifications faster than unmodified tRNA, and DusA may exhibit an even stronger preference for modifying an already modified tRNA substrate that it does for tRNA binding. *T. thermophilus*, *S. cerevisiae*, and human U20-dihydrouridylating enzymes have all shown either strong preferences or absolute requirements for modifications in their tRNA substrate for efficient binding and/or modification. Similarly, modifications have been suggested to be a requirement for *Tth*DusA•tRNA complex formation (32) and/or increase reaction velocities at high temperatures (38). Whereas purified yeast Dus2 has dihydrouridine activity on *in vitro* transcribed precursor (pre)-tRNA^Tyr^ and pre-tRNA^Leu^ (59), the affinity of yeast Dus2 for native mature tRNA^Leu^ lacking D20 was stronger than that for the pre-tRNA transcript, and the oxidation of yeast Dus2 is >600-fold faster in the presence of native, modified tRNA^Leu^ compared to an *in vitro* transcript (36). Finally, human Dus2 has been shown to bind an unmodified tRNA^Asp^ transcript but no dihydrouridine formation was observed for this unmodified substrate tRNA *in vitro* (37), suggesting modifications are required for dihydrouridylation activity.

These findings suggest D20 is added to tRNA in the later stages of tRNA maturation. However, D20 is formed prior to intron removal in *Xenopus* oocytes and yeast indicating further modifications are added following D20 onto spliced tRNA (59–61). Further to this, we observe a lower proportion of tRNAs contain acp^3^U47 in *E. coli* lacking *dusA*, implying TapT may modify U47 at an even later stage of tRNA maturation. In conclusion, the introduction of D20 seems to occur in the intermediate stages during tRNA maturation across bacteria and eukaryotes.

### DusA binds folded tRNA using a two-step mechanism involving a local conformational change in the D arm

Our kinetic experiments reveal that DusA binds tRNA with a two-step mechanism including a tRNA conformational change. Upon *Tth*Dus binding to *Tth* tRNA^Phe^, significant conformational changes are observed for U16, U17, and the target base U20, but G18 and G19 remain stably base paired with U55 and C56 in the T arm, respectively, and there are no significant tRNA conformational changes outside of the D loop (32). Moreover, tRNA variants with disrupted G18-U55 and/or G19-C56 interactions cannot be modified by yeast Dus2 (58). Therefore, we speculate that the conformational change we observe corresponds to these local conformational rearrangements within the D loop changing the environment of the fluorophore attached at U8).

This local conformational change within the D loop upon DusA binding contrasts the major rearrangements within the tRNA elbow that must take place for TrmA and TruB to bind their respective target bases, U54 and U55. Upon binding TrmA, U54 flips out of the T loop leading to a base stack containing G53, A58, G57, C56, and U55 and disruption of the G18-U55 and G19-C56 base pairs (62). Similarly, U55, C56, and G57 flip out of the T loop upon binding TruB, again disrupting tertiary base pairing between these nucleobases and G18 and G19 in the D arm (63,64). Accordingly, partially unfolded tRNA substrates with disrupted G18-U55 and/or G19-C56 interactions are modified by TrmA with a higher catalytic efficiency (65), and display faster T arm base flipping kinetics upon interaction with TruB (34). Taken together, these structural and biochemical experiments reveal a fundamental mechanistic difference between DusA and TrmA/TruB: whereas TrmA and TruB enzymes must break tertiary interactions in the tRNA elbow to access their substrate, DusA instead requires a stable elbow structure in order to modify tRNAs.

### DusA has distinct cellular functions from TrmA and TruB

As discussed above, D20 synthases like DusA show a conserved preference for acting in the intermediate stages of tRNA maturation acting on folded tRNA whereas the tRNA chaperones TrmA and TruB prefer to act in the early stages where they unfold tRNA and provide a second chance at correct tRNA folding. These differences in molecular mechanisms between DusA and TrmA/TruB result in distinct cellular functions for these enzymes. Previously, we have shown that tRNA chaperones TrmA and TruB enhance aminoacylation globally across all tRNA species in *E. coli* while not affecting steady-state tRNA abundances (20). In contrast, we discover here that deletion of *dusA* does not affect global aminoacylation but the charging levels of two tRNA^Gly^ isoacceptors are significantly reduced. Although the aminoacylation of tRNA^Gly^_UCC_ is decreased, this tRNA has not been previously shown to contain U20/U20a (18). These observations may suggest a more specific and perhaps indirect role for DusA in modulating the aminoacylation and possibly abundance of tRNA^Gly^ species without broadly affecting the charging or cellular stability of other tRNAs. Intriguingly, previous work has shown that upon oxidative stress, the cellular aminoacylation level is significantly reduced for all three Gly isoacceptors (52). Since DusA utilizes reducing equivalents, deletion of *dusA* may affect cellular redox homeostasis and/or play a role underlaying the loss of active tRNA^Gly^ in oxidative stress; however, these processes require further study. Except for tRNA^Gly^ isoaccpetors, almost every other tRNA has a higher abundance in the *dusA* knockout indicating on a global scale, tRNA abundance is increased in the *ΔdusA* strain. The biological relevance for these increased tRNA levels remains unclear but may reflect compensation for hypomodified tRNAs or changes in cellular redox homeostasis. Intriguingly, when DusA is overexpressed in *E. coli*, cells display a noticeably yellow phenotype (data not shown), suggesting that FMN production is upregulated when DusA is overexpressed by so far unknown mechanisms.

Despite minimal changes in tRNA abundance and aminoacylation upon *dusA* deletion, we observe decreases in translation for several codons irrespective of the changes in tRNA abundance and/or tRNA aminoacylation. In contrast to the deletion of *trmA* and/or *truB* which cause both decreases and increases in codon-specific translation, deletion of *dusA* consistently negatively impacts the translation of specific codons or does not impact the translation of a given codon at all. A recent study has revealed the dihydrouridine level of tRNA does not affect tRNA recruitment to polysomes in *E. coli* (18), suggesting tRNA binding by EF-Tu is likely not impaired in the absence of *dusA*. Thus, dihydrouridylation of tRNAs by DusA must directly impact the function of specific tRNAs on the ribosome during mRNA translation for specific codons. Related to a proposed function of m^5^U54 formed by TrmA (22), we speculate that D20 affects the flexibility of tRNA required to move through the ribosome during translocation from the A to the P site. Further supporting a role for U20 modification for translation, addition of purified human Dus2 to a rabbit reticulocyte *in vitro* translation system increased synthesis of certain proteins (26).

In conclusion, our results suggest that DusA acts non-redundantly with TrmA and TruB to fine-tune tRNA function during mRNA translation (Figure 8). Whereas TrmA and TruB disrupt interactions between the D and T arms to modify all tRNAs thereby providing all tRNAs a second chance at properly folding, DusA instead requires the tRNA elbow to be already folded properly prior to dihydrouridylation of specific tRNAs. As tRNA modifications are known to stabilize the tRNA structure (5) and *Tth*Dus does not form any specific interactions with tRNA bases that undergo modification (32), DusA likely prefers to bind an already modified tRNA substrate due to the reinforced tertiary structure. Because TrmA and TruB disrupt the tRNA tertiary structure, these enzymes provide misfolded tRNAs second chances at properly folding (34,35). In contrast, DusA is unlikely to be a tRNA chaperone as it requires folded tRNA which retains its tertiary structure when interacting with DusA. Accordingly, TrmA and TruB are known to act early in tRNA maturation (40,66,67), whereas DusA most likely acts in the intermediate or late stages. In line with the role of TrmA and TruB as tRNA chaperones, these enzymes increase aminoacylation of all tRNAs (20), while DusA does not affect tRNA charging on a global scale. Like for TrmA and TruB, many changes in codon specific translation are observed in the absence of DusA in *E. coli*; however, several codon-specific changes for TrmA and TruB may be accounted for by aminoacylation changes, whereas most codon-specific alterations in the *dusA* knockout strain can only be explained by differences in tRNA activity on the ribosome. Since DusC also does not disrupt tertiary interactions in the tRNA elbow (68), we speculate that DusC likewise may not function as a tRNA chaperone and instead acts similarly to DusA in tRNA maturation. In conclusion, sophisticated *in vitro* studies combined with global *in vivo* analyses reveal the diverse and complementary functions of different tRNA modification enzymes that collectively fine-tune tRNAs for optimal protein synthesis.

**Figure 8.**
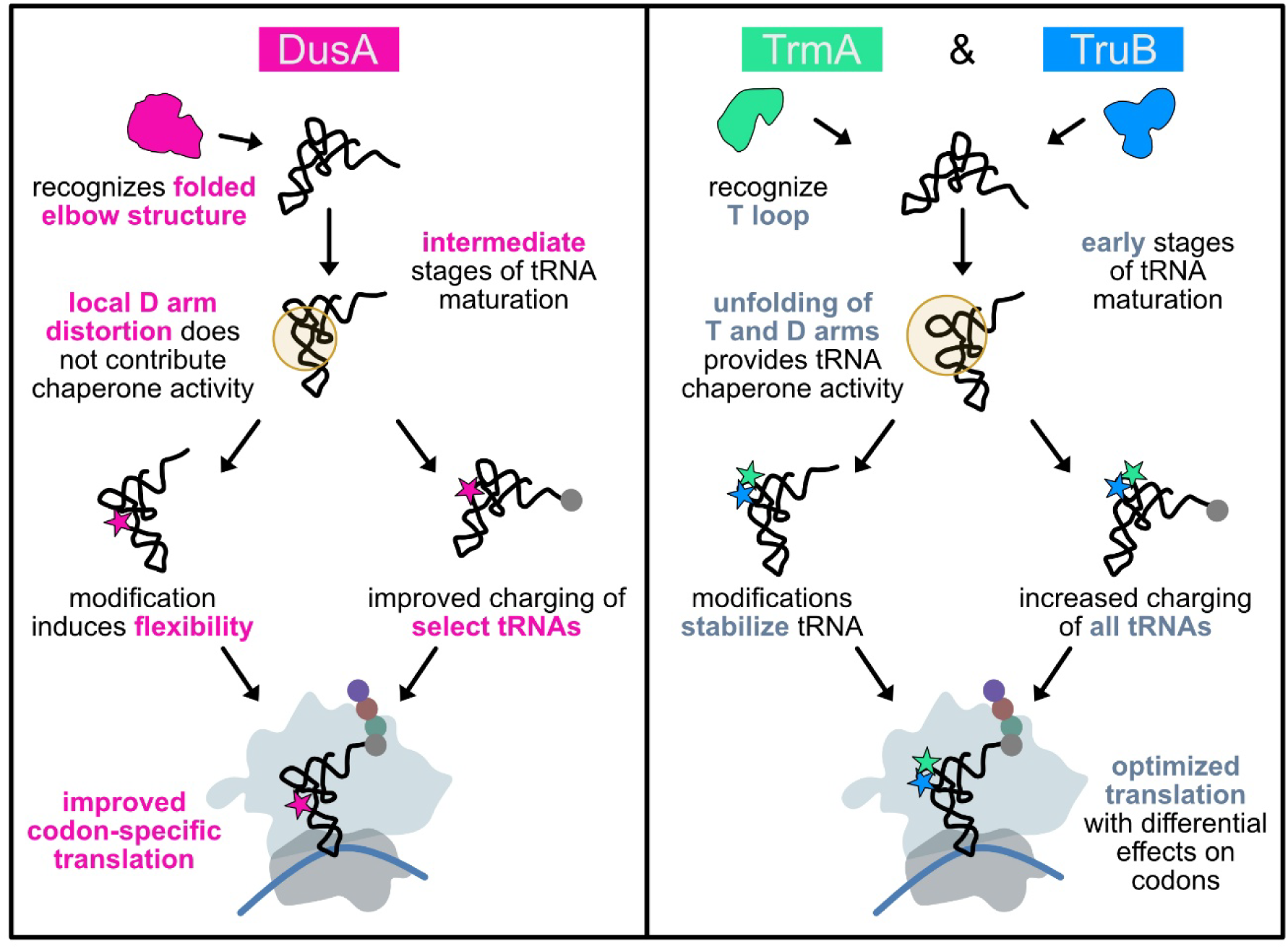
Comparison of DusA with T arm modifying enzymes TrmA and TruB. Although all three enzymes modify the tRNA elbow region, DusA differs markedly in its molecular mechanism and biological function.

## Acknowledgements

We thank Assaf Katz for the kind gift of the sfGFP codon reporter library, Kevin Li for purifying DusA variants, Saskia Funk for initial stopped-flow experiments, and Timothy Vos for sub-cloning the pET28a-DusA vector. *E. coli* BW25113 wildtype and *ΔdusA* Keio collection strains were obtained from National BioResource Project (National Institute of Genetics, Japan).

## Author contributions

S.K.S. and U.K. conceptualized the study. N.H. performed and analyzed variant stopped-flow experiments and certain codon reporter assays. L.B. performed and analyzed mass spectrometry experiments. S.K.S. performed all other experiments and data analysis. S.K.S. wrote the first draft of the manuscript and U.K. and S.K.S. edited the manuscript. U.K. and K.S.K. provided funding and supervised the project.

## Data availability

MSR-seq data is available at the NCBI GEO database under the accession GSE304803.

## References

1. Phizicky, E.M. and Hopper, A.K. (2023) The life and times of a tRNA. RNA, 29, 898–957.

2. Schultz, S.K. and Kothe, U. (2024) RNA modifying enzymes shape tRNA biogenesis and function. J Biol Chem, 300, 107488.

3. Brégeon, D., Pecqueur, L., Toubdji, S., Sudol, C., Lombard, M., Fontecave, M., de Crécy-Lagard, V., Motorin, Y., Helm, M. and Hamdane, D. (2022) Dihydrouridine in the Transcriptome: New Life for This Ancient RNA Chemical Modification. ACS Chem Biol, 17, 1638–1657.

4. Finet, O., Yague-Sanz, C., Marchand, F. and Hermand, D. (2022) The Dihydrouridine landscape from tRNA to mRNA: a perspective on synthesis, structural impact and function. RNA Biol, 19, 735–750.

5. Lorenz, C., Lünse, C.E. and Mörl, M. (2017) tRNA Modifications: Impact on Structure and Thermal Adaptation. Biomolecules, 7.

6. Dalluge, J.J., Hashizume, T., Sopchik, A.E., McCloskey, J.A. and Davis, D.R. (1996) Conformational flexibility in RNA: the role of dihydrouridine. Nucleic Acids Res, 24, 1073–1079.

7. Dyubankova, N., Sochacka, E., Kraszewska, K., Nawrot, B., Herdewijn, P. and Lescrinier, E. (2015) Contribution of dihydrouridine in folding of the D-arm in tRNA. Org Biomol Chem, 13, 4960–4966.

8. Nomura, Y., Ohno, S., Nishikawa, K. and Yokogawa, T. (2016) Correlation between the stability of tRNA tertiary structure and the catalytic efficiency of a tRNA-modifying enzyme, archaeal tRNA-guanine transglycosylase. Genes Cells, 21, 41–52.

9. Dalluge, J.J., Hamamoto, T., Horikoshi, K., Morita, R.Y., Stetter, K.O. and McCloskey, J.A. (1997) Posttranscriptional modification of tRNA in psychrophilic bacteria. J Bacteriol, 179, 1918–1923.

10. Noon, K.R., Guymon, R., Crain, P.F., McCloskey, J.A., Thomm, M., Lim, J. and Cavicchioli, R. (2003) Influence of temperature on tRNA modification in archaea: Methanococcoides burtonii (optimum growth temperature [Topt], 23 degrees C) and Stetteria hydrogenophila (Topt, 95 degrees C). J Bacteriol, 185, 5483–5490.

11. Kasprzak, J.M., Czerwoniec, A. and Bujnicki, J.M. (2012) Molecular evolution of dihydrouridine synthases. BMC Bioinformatics, 13, 153.

12. Bou-Nader, C., Montémont, H., Guérineau, V., Jean-Jean, O., Brégeon, D. and Hamdane, D. (2018) Unveiling structural and functional divergences of bacterial tRNA dihydrouridine synthases: perspectives on the evolution scenario. Nucleic Acids Res, 46, 1386–1394.

13. Bishop, A.C., Xu, J., Johnson, R.C., Schimmel, P. and de Crécy-Lagard, V. (2002) Identification of the tRNA-dihydrouridine synthase family. J Biol Chem, 277, 25090–25095.

14. Toubdji, S., Thullier, Q., Kilz, L.M., Marchand, V., Yuan, Y., Sudol, C., Goyenvalle, C., Jean-Jean, O., Rose, S., Douthwaite, S. et al. (2024) Exploring a unique class of flavoenzymes: Identification and biochemical characterization of ribosomal RNA dihydrouridine synthase. Proc Natl Acad Sci U S A, 121, e2401981121.

15. Faivre, B., Lombard, M., Fakroun, S., Vo, C.D., Goyenvalle, C., Guérineau, V., Pecqueur, L., Fontecave, M., De Crécy-Lagard, V., Brégeon, D. and Hamdane, D. (2021) Dihydrouridine synthesis in tRNAs is under reductive evolution in Mollicutes. RNA Biol, 18, 2278–2289.

16. Sudol, C., Kilz, L.M., Marchand, V., Thullier, Q., Guérineau, V., Goyenvalle, C., Faivre, B., Toubdji, S., Lombard, M., Jean-Jean, O. et al. (2024) Functional redundancy in tRNA dihydrouridylation. Nucleic Acids Res, 52, 5880–5894.

17. Xing, F., Hiley, S.L., Hughes, T.R. and Phizicky, E.M. (2004) The specificities of four yeast dihydrouridine synthases for cytoplasmic tRNAs. J Biol Chem, 279, 17850–17860.

18. Kilz, L.M., Zimmermann, S., Marchand, V., Bourguignon, V., Sudol, C., Brégeon, D., Hamdane, D., Motorin, Y. and Helm, M. (2024) Differential redox sensitivity of tRNA dihydrouridylation. Nucleic Acids Res, 52, 12784–12797.

19. Farrugia, D.N., Elbourne, L.D., Mabbutt, B.C. and Paulsen, I.T. (2015) A novel family of integrases associated with prophages and genomic islands integrated within the tRNA-dihydrouridine synthase A (dusA) gene. Nucleic Acids Res, 43, 4547–4557.

20. Schultz, S.K., Katanski, C.D., Halucha, M., Peña, N., Fahlman, R.P., Pan, T. and Kothe, U. (2024) Modifications in the T arm of tRNA globally determine tRNA maturation, function, and cellular fitness. Proc Natl Acad Sci U S A, 121, e2401154121.

21. Babosan, A., Fruchard, L., Krin, E., Carvalho, A., Mazel, D. and Baharoglu, Z. (2022) Nonessential tRNA and rRNA modifications impact the bacterial response to sub-MIC antibiotic stress. Microlife, 3, uqac019.

22. Jones, J.D., Franco, M.K., Tardu, M., Smith, T.J., Snyder, L.R., Eyler, D.E., Polikanov, Y., Kennedy, R.T., Niederer, R.O. and Koutmou, K.S. (2023) Conserved 5-methyluridine tRNA modification modulates ribosome translocation. bioRxiv.

23. Durand, R., Deschênes, F. and Burrus, V. (2021) Genomic islands targeting dusA in Vibrio species are distantly related to Salmonella Genomic Island 1 and mobilizable by IncC conjugative plasmids. PLoS Genet, 17, e1009669.

24. Tan, X., Zhang, M., Liu, S., Xiao, X., Zhang, Y. and Jian, H. (2024) Prophage enhances the ability of deep-sea bacterium Shewanella psychrophila WP2 to utilize D-amino acid. Microbiol Spectr, 12, e0326323.

25. Kuchino, Y. and Borek, E. (1978) Tumour-specific phenylalanine tRNA contains two supernumerary methylated bases. Nature, 271, 126–129.

26. Kato, T., Daigo, Y., Hayama, S., Ishikawa, N., Yamabuki, T., Ito, T., Miyamoto, M., Kondo, S. and Nakamura, Y. (2005) A novel human tRNA-dihydrouridine synthase involved in pulmonary carcinogenesis. Cancer Res, 65, 5638–5646.

27. Mittelstadt, M., Frump, A., Khuu, T., Fowlkes, V., Handy, I., Patel, C.V. and Patel, R.C. (2008) Interaction of human tRNA-dihydrouridine synthase-2 with interferon-induced protein kinase PKR. Nucleic Acids Res, 36, 998–1008.

28. Chen, X., Ji, B., Hao, X., Li, X., Eisele, F., Nyström, T. and Petranovic, D. (2020) FMN reduces Amyloid-β toxicity in yeast by regulating redox status and cellular metabolism. Nat Commun, 11, 867.

29. Li, Z., Yin, C., Li, B., Yu, Q.Y., Mao, W.J., Li, J., Lin, J.P., Meng, Y.Q., Feng, H.M. and Jing, T. (2020) DUS4L Silencing Suppresses Cell Proliferation and Promotes Apoptosis in Human Lung Adenocarcinoma Cell Line A549. Cancer Manag Res, 12, 9905–9913.

30. Draycott, A.S., Schaening-Burgos, C., Rojas-Duran, M.F., Wilson, L., Schärfen, L., Neugebauer, K.M., Nachtergaele, S. and Gilbert, W.V. (2022) Transcriptome-wide mapping reveals a diverse dihydrouridine landscape including mRNA. PLoS Biol, 20, e3001622.

31. Finet, O., Yague-Sanz, C., Krüger, L.K., Tran, P., Migeot, V., Louski, M., Nevers, A., Rougemaille, M., Sun, J., Ernst, F.G.M. et al. (2022) Transcription-wide mapping of dihydrouridine reveals that mRNA dihydrouridylation is required for meiotic chromosome segregation. Mol Cell, 82, 404–419.e409.

32. Yu, F., Tanaka, Y., Yamashita, K., Suzuki, T., Nakamura, A., Hirano, N., Suzuki, T., Yao, M. and Tanaka, I. (2011) Molecular basis of dihydrouridine formation on tRNA. Proc Natl Acad Sci U S A, 108, 19593–19598.

33. Savage, D.F., de Crécy-Lagard, V. and Bishop, A.C. (2006) Molecular determinants of dihydrouridine synthase activity. FEBS Lett, 580, 5198–5202.

34. Keffer-Wilkes, L.C., Veerareddygari, G.R. and Kothe, U. (2016) RNA modification enzyme TruB is a tRNA chaperone. Proc Natl Acad Sci U S A, 113, 14306–14311.

35. Keffer-Wilkes, L.C., Soon, E.F. and Kothe, U. (2020) The methyltransferase TrmA facilitates tRNA folding through interaction with its RNA-binding domain. Nucleic Acids Res, 48, 7981–7990.

36. Rider, L.W., Ottosen, M.B., Gattis, S.G. and Palfey, B.A. (2009) Mechanism of dihydrouridine synthase 2 from yeast and the importance of modifications for efficient tRNA reduction. J Biol Chem, 284, 10324–10333.

37. Bou-Nader, C., Pecqueur, L., Bregeon, D., Kamah, A., Guérineau, V., Golinelli-Pimpaneau, B., Guimarães, B.G., Fontecave, M. and Hamdane, D. (2015) An extended dsRBD is required for post-transcriptional modification in human tRNAs. Nucleic Acids Res, 43, 9446–9456.

38. Kusuba, H., Yoshida, T., Iwasaki, E., Awai, T., Kazayama, A., Hirata, A., Tomikawa, C., Yamagami, R. and Hori, H. (2015) In vitro dihydrouridine formation by tRNA dihydrouridine synthase from Thermus thermophilus, an extreme-thermophilic eubacterium. J Biochem, 158, 513–521.

39. Wright, J.R., Keffer-Wilkes, L.C., Dobing, S.R. and Kothe, U. (2011) Pre-steady-state kinetic analysis of the three Escherichia coli pseudouridine synthases TruB, TruA, and RluA reveals uniformly slow catalysis. Rna, 17, 2074–2084.

40. Schultz, S.K. and Kothe, U. (2020) tRNA elbow modifications affect the tRNA pseudouridine synthase TruB and the methyltransferase TrmA. RNA, 26, 1131–1142.

41. Gill, S.C. and von Hippel, P.H. (1989) Calculation of protein extinction coefficients from amino acid sequence data. Anal Biochem, 182, 319–326.

42. Sampson, J.R., DiRenzo, A.B., Behlen, L.S. and Uhlenbeck, O.C. (1989) Nucleotides in yeast tRNAPhe required for the specific recognition by its cognate synthetase. Science, 243, 1363–1366.

43. Peterson, E.T. and Uhlenbeck, O.C. (1992) Determination of recognition nucleotides for Escherichia coli phenylalanyl-tRNA synthetase. Biochemistry, 31, 10380–10389.

44. Schultz, S.K. and Kothe, U. (2021) Partially modified tRNAs for the study of tRNA maturation and function. Methods Enzymol, 658, 225–250.

45. Jones, J.D., Franco, M.K., Smith, T.J., Snyder, L.R., Anders, A.G., Ruotolo, B.T., Kennedy, R.T. and Koutmou, K.S. (2023) Methylated guanosine and uridine modifications in S. cerevisiae mRNAs modulate translation elongation. RSC Chem Biol, 4, 363–378.

46. Schultz, S.K. and Kothe, U. (2023) Fluorescent labeling of tRNA for rapid kinetic interaction studies with tRNA-binding proteins. Methods Enzymol, 692, 103–126.

47. Schultz, S.K., Meadows, K. and Kothe, U. (2023) Molecular mechanism of tRNA binding by the Escherichia coli N7 guanosine methyltransferase TrmB. J Biol Chem, 299, 104612.

48. Baba, T., Ara, T., Hasegawa, M., Takai, Y., Okumura, Y., Baba, M., Datsenko, K.A., Tomita, M., Wanner, B.L. and Mori, H. (2006) Construction of Escherichia coli K-12 in-frame, single-gene knockout mutants: the Keio collection. Mol Syst Biol, 2, 2006.0008.

49. Watkins, C.P., Zhang, W., Wylder, A.C., Katanski, C.D. and Pan, T. (2022) A multiplex platform for small RNA sequencing elucidates multifaceted tRNA stress response and translational regulation. Nat Commun, 13, 2491.

50. Chan, P.P. and Lowe, T.M. (2016) GtRNAdb 2.0: an expanded database of transfer RNA genes identified in complete and draft genomes. Nucleic Acids Res, 44, D184–189.

51. Langmead, B. and Salzberg, S.L. (2012) Fast gapped-read alignment with Bowtie 2. Nat Methods, 9, 357–359.

52. Leiva, L.E., Elgamal, S., Leidel, S.A., Orellana, O., Ibba, M. and Katz, A. (2022) Oxidative stress strongly restricts the effect of codon choice on the efficiency of protein synthesis in Escherichia coli. Front Microbiol, 13, 1042675.

53. Chen, X., Sim, S., Wurtmann, E.J., Feke, A. and Wolin, S.L. (2014) Bacterial noncoding Y RNAs are widespread and mimic tRNAs. Rna, 20, 1715–1724.

54. Boccaletto, P., Stefaniak, F., Ray, A., Cappannini, A., Mukherjee, S., Purta, E., Kurkowska, M., Shirvanizadeh, N., Destefanis, E., Groza, P. et al. (2022) MODOMICS: a database of RNA modification pathways. 2021 update. Nucleic Acids Res, 50, D231–d235.

55. Meyer, B., Immer, C., Kaiser, S., Sharma, S., Yang, J., Watzinger, P., Weiss, L., Kotter, A., Helm, M., Seitz, H.M. et al. (2020) Identification of the 3-amino-3-carboxypropyl (acp) transferase enzyme responsible for acp^3^U formation at position 47 in *Escherichia coli* tRNAs. Nucleic Acids Res, 48, 1435–1450.

56. Takakura, M., Ishiguro, K., Akichika, S., Miyauchi, K. and Suzuki, T. (2019) Biogenesis and functions of aminocarboxypropyluridine in tRNA. Nat Commun, 10, 5542.

57. Byrne, R.T., Konevega, A.L., Rodnina, M.V. and Antson, A.A. (2010) The crystal structure of unmodified tRNAPhe from Escherichia coli. Nucleic Acids Res, 38, 4154–4162.

58. Cavaillé, J., Chetouani, F. and Bachellerie, J.P. (1999) The yeast Saccharomyces cerevisiae YDL112w ORF encodes the putative 2’-O-ribose methyltransferase catalyzing the formation of Gm18 in tRNAs. Rna, 5, 66–81.

59. Xing, F., Martzen, M.R. and Phizicky, E.M. (2002) A conserved family of Saccharomyces cerevisiae synthases effects dihydrouridine modification of tRNA. Rna, 8, 370–381.

60. Nishikura, K. and De Robertis, E.M. (1981) RNA processing in microinjected Xenopus oocytes. Sequential addition of base modifications in the spliced transfer RNA. J Mol Biol, 145, 405–420.

61. Etcheverry, T., Colby, D. and Guthrie, C. (1979) A precursor to a minor species of yeast tRNASer contains an intervening sequence. Cell, 18, 11–26.

62. Alian, A., Lee, T.T., Griner, S.L., Stroud, R.M. and Finer-Moore, J. (2008) Structure of a TrmA-RNA complex: A consensus RNA fold contributes to substrate selectivity and catalysis in m5U methyltransferases. Proc Natl Acad Sci U S A, 105, 6876–6881.

63. Pan, H., Agarwalla, S., Moustakas, D.T., Finer-Moore, J. and Stroud, R.M. (2003) Structure of tRNA pseudouridine synthase TruB and its RNA complex: RNA recognition through a combination of rigid docking and induced fit. Proc Natl Acad Sci U S A, 100, 12648–12653.

64. Hoang, C. and Ferré-D’Amaré, A.R. (2001) Cocrystal structure of a tRNA Psi55 pseudouridine synthase: nucleotide flipping by an RNA-modifying enzyme. Cell, 107, 929–939.

65. Kealey, J.T., Gu, X. and Santi, D.V. (1994) Enzymatic mechanism of tRNA (m5U54)methyltransferase. Biochimie, 76, 1133–1142.

66. Barraud, P., Gato, A., Heiss, M., Catala, M., Kellner, S. and Tisne, C. (2019) Time-resolved NMR monitoring of tRNA maturation. Nat Commun, 10, 3373.

67. Heiss, M., Hagelskamp, F., Marchand, V., Motorin, Y. and Kellner, S. (2021) Cell culture NAIL-MS allows insight into human tRNA and rRNA modification dynamics in vivo. Nat Commun, 12, 389.

68. Byrne, R.T., Jenkins, H.T., Peters, D.T., Whelan, F., Stowell, J., Aziz, N., Kasatsky, P., Rodnina, M.V., Koonin, E.V., Konevega, A.L. and Antson, A.A. (2015) Major reorientation of tRNA substrates defines specificity of dihydrouridine synthases. Proc Natl Acad Sci U S A, 112, 6033–6037.

